# Strain-dependent induction of primary bile acid 7-dehydroxylation by cholic acid

**DOI:** 10.1101/2022.02.15.480494

**Authors:** Eduard Vico-Oton, Colin Volet, Nicolas Jacquemin, Yuan Dong, Siegfried Hapfelmeier, Karin Lederballe Meibom, Rizlan Bernier-Latmani

## Abstract

Bile acids (BAs) are steroid-derived molecules with important roles in digestion, the maintenance of host metabolism and immunomodulation. Primary BAs are synthesised by the host, while secondary BAs are produced by the gut microbiome through transformation of the former. Regulation of microbial production of secondary BAs is not well understood, particularly the production of 7-dehydroxylated BAs, which are the most potent agonists for host BA receptors. The 7-dehydroxylation of cholic acid (CA) is well established and is linked to the expression of a bile acid-inducible (*bai*) operon responsible for this process. However, little to no 7-dehydroxylation has been reported for other host-derived BAs (e.g., chenodeoxycholic acid, CDCA or ursodeoxycholic acid, UDCA). Here, we demonstrate that the 7-dehydroxylation of CDCA and UDCA by *Clostridium scindens* is induced by CA suggesting that CA-dependent transcriptional regulation of 7-dehydroxylation is generalisable to CDCA and UDCA. In contrast, the murine isolate *Extibacter muris* did not respond to CA exposure *in vitro*, suggesting that *bai* genes are regulated differently in this strain. However, it could 7-dehydroxylate *in vivo* and its *in vitro* activity was promoted by the addition of cecal content. The accessory gene *baiJ* was only upregulated in the *Clostridium scindens* ATCC 35704 strain, implying mechanistic differences amongst isolates. Interestingly, the human-derived *C. scindens* strains were also capable of 7-dehydroxylating murine bile acids (muricholic acids) to a limited extent. This study shows novel 7-dehydroxylation activity *in vitro* as a result of CA-driven induction and suggests distinct *bai* gene induction mechanisms across bacterial species.

## Introduction

Primary bile acids (BAs) are metabolites synthesised from cholesterol by hepatocytes while secondary BAs are produced by the gut microbiome through the transformation of primary BAs (Figure 1). In the liver, the BA are conjugated to glycine or taurine. The three main microbial BA transformations are deconjugation (loss of the amino acid group), oxidation (of one or several of the hydroxyl groups), and 7α-dehydroxylation (7-DH-ion), the loss of a hydroxyl group at the C7 position^1^. These microbial transformations increase the diversity of the BA pool (Figure 1) and enhance BA affinity to host receptors. In particular, 7-DH-ion turns primary BAs such as cholic acid (CA) and chenodeoxycholic acid (CDCA) into the 7-dehydroxylated (7-DH-ed) BAs deoxycholic acid (DCA) and lithocholic acid (LCA), respectively^2^.

**Figure 1.**
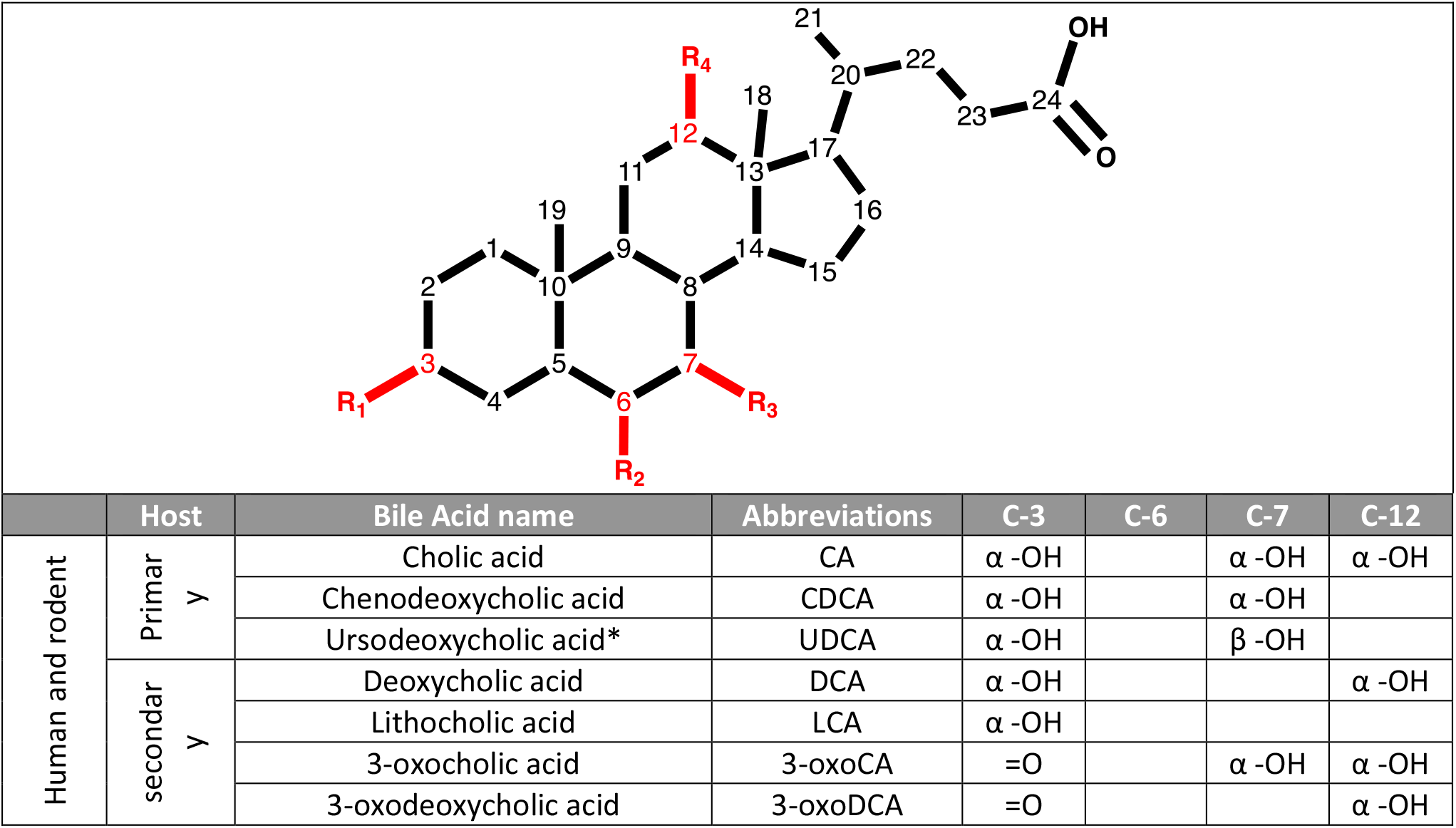
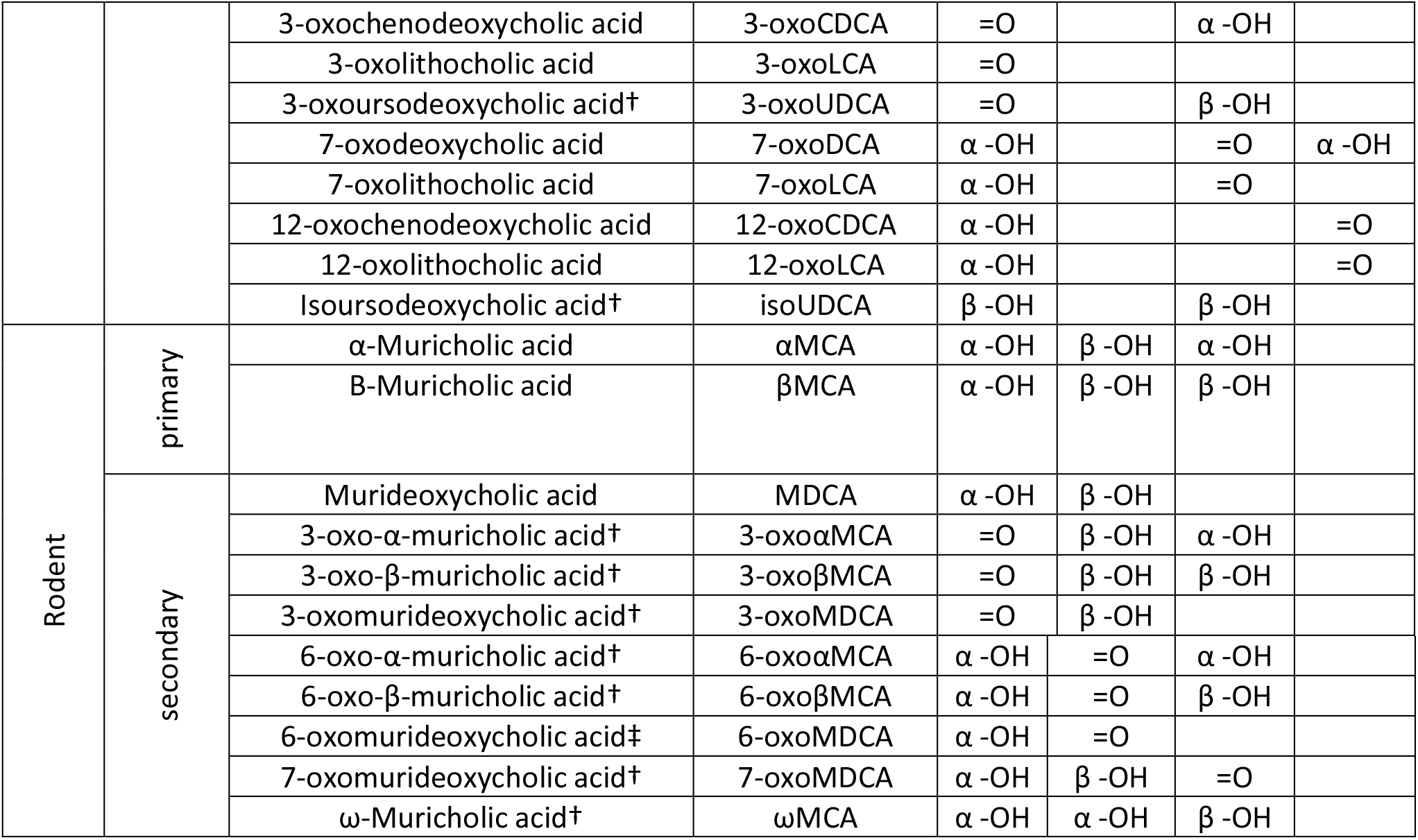
List of deconjugated human and rodent bile acids (BAs) discussed here. The characteristic that distinguishes BAs is the presence of a hydroxyl group at the C-3, C-6, C-7 and/or C-12 position. The hydroxyl groups can be in α-or β-conformation, oxidised into a ketone group, or fully removed (dehydroxylated BA). CA and CDCA are primary BAs of both humans and rodents whereas MCAs are exclusively produced by rodents. (*) UDCA is a primary BA in rodents while it is a secondary BA in humans where the gut microbes epimerise it from CDCA. (†) Bile acids that might have been detected in this study but for which no standards were available. (‡) Also known as 6-oxolithocholic acid (6-oxoLCA). See Supplementary Table 3 for full chemical names of the BAs.

BAs act as detergents to solubilise dietary fats, but also have important metabolic and immunomodulatory roles through activation of their target receptors^3^. The two best-studied BA receptors are the Farnesoid X Receptor (FXR), a nuclear receptor, and the G Protein-Coupled Bile Acid Receptor, GPBAR1, also known as Takeda G-Protein Receptor 5 (TGR5), which is a membrane receptor. FXR is activated through the binding of BA agonists, particularly the 7-DH-ed LCA but also DCA^4^. FXR activation results in the inhibition of primary BA synthesis through repression of the cholesterol 7α-hydroxylase CYP7A1. Regulation of BA production limits BA concentration and therefore, toxicity. On the other hand, BA dysregulation can cause health issues such as cholestasis, irritable bowel syndrome, gallstone disease, or even the induction of colorectal cancer^5–7^. Besides BA homeostasis, FXR also has focal roles in glucose and lipid homeostasis^8^. Similarly, TGR5 is a multifunctional regulator involved in glucose homeostasis, energy expenditure, and the modulation of the inflammatory response^9,10^. LCA, DCA and their tauro-conjugated forms TLCA and TDCA, are among the strongest agonists of TGR5^11,12^, highlighting the importance of microbial transformation, particularly 7-DH-ion, in TGR5 activation. Moreover, DCA and LCA have protective properties against *Clostridium difficile* infection^13,14^.

The study of BAs has traditionally been based on mouse models^15^. Besides CA and CDCA, mice (and other rodents) also generate muricholic acids (MCAs) such as α-MCA and β-MCA (Figure 1) and re-hydroxylate DCA and LCA in the liver^16^. Additionally, the mouse liver is capable of producing primary ursodeoxycholic acid (UDCA)^16^ although there is evidence that the gut microbiome is responsible for a significant fraction of UDCA in the gut^14^. On the other hand, UDCA is exclusively a secondary BA in humans^7^. Because 7-DH-ion plays a major role in host homeostasis, significant effort has been expended to study 7-dehydroxylating (7-DH-ing) bacteria.

Nonetheless, experimental evidence of 7-DH-ion is limited to a few species of the *Clostridiales* order. One of the best characterised is the human isolate *Clostridium scindens* ATCC 35704, the type strain of *C. scindens*^17^.The *Extibacter muris* DSM 28560 (JM40) strain was recently isolated from mice and identified as a 7-DH-ing organism^18,19^. The ability of *C. scindens* ATCC 35704 to 7-dehydroxylate both *in vivo* and *in vitro* is well established^20^; *E. muris* has also been shown to 7-dehydroxylate *in vivo*, transforming primary BAs CA, CDCA, αMCA, βMCA and UDCA into their respective secondary BAs DCA, LCA and MDCA. Previous research had focused on *E. muris* strain DSM 28560^21^ and in this study, we demonstrate that *E. muris* strain DSM 28561 (SJ24) also has the ability to 7-dehydroxylate (7-DH-ate) *in vivo*.

The biochemical machinery for 7-DH-ion is encoded in the *bai* (bile acid inducible) eight-gene operon (*baiBCDEA2FGHI*)^22^. In addition, the *C. scindens* ATCC 35704 strain harbours the accessory gene *baiJ* (HDCHBGLK_03451)^23^ whereas the *E. muris* DSM 28650 genome includes a *baiJKL* pseudogene cluster^21^. Most of the published work on the 7-DH-ion pathway has been performed with another *C. scindens* strain, VPI 12708^24,25^. A recent publication by Funabashi *et al*. cloned the *bai* operon of *C. scindens* VPI 12708 into *Clostridium sporogenes* which then showed *in vivo* 7-DH-ing activity^25^. Different strains exhibit varying efficiency in 7-DH-ing CA *in vitro*. The *C. scindens* ATCC 35704 and VPI 12708 strains show rapid transformation to DCA while *E. muris* DSM 28560 has more limited activity^19,20,26^. Other known 7-DH-ing strains such as *Clostridium hylemonae* and *Peptacetobacter hiranonis* have been reported as harbouring weak and strong activity, respectively^27^ and a new strain of *P. hiranonis* recently isolated from dog faeces displayed *in vitro* 7-DH-ion at around 30% conversion of CA to DCA^28^. Notably, *in vitro* 7-DH-ion of other primary BAs has been reported to be minor (CDCA) or non-existent (MCAs and UDCA)^20,21,29^.

The limited *in vitro* 7-DH-ion of primary BAs other than CA (i.e., CDCA and MCAs) is striking considering that secondary 7-DH-ed forms of these BAs are routinely detected at significant concentrations in the host^30,31^. Most studies tackling *in vitro* primary bile acid 7-DH-ion consider each BA in isolation. In addition, significant overexpression of the *bai* operon in response to CA has been reported^32,33^ but there is no information about the potential induction of this operon by other primary BAs (i.e., CDCA and MCAs) and whether induction by CA also results in the transformation of the latter. We hypothesise that CA-dependent induction of the *bai* operon promotes 7-DH-ing activity of other BAs when they occur together with CA. Moreover, we posit that primary BAs other than CA cannot induce their own transformation.

Here, the expression of *bai* genes was measured *in vitro* in the presence of CA, CDCA, αMCA, βMCA and UDCA, with and without co-induction with CA to test whether the overexpression of *bai* genes was exclusive to CA and whether CA-induced overexpression was sufficient to promote the 7-DH-ion of other BAs. The experiments were performed with three strains, the human isolates *C. scindens* ATCC 35704 and VPI 12708 and the murine isolate *E. muris* DSM 28561 (SJ24). The results show that the response to CA co-induction was strain-variable. It was highly effective for *C. scindens* strains and sufficient to promote the transformation of other primary BAs. For *E. muris*, none of the BAs tested promoted 7-DH-ion, nevertheless, a positive effect was observed when the bacterium was co-cultured with a small amount of faecal content from germ-free mice, suggesting that signaling from the host may be responsible for the induction of 7-DH-ion in *E. muris* SJ24.

This work highlights the importance of the presence CA for the 7-DH-ion of other BAs. Moreover, results from *E. muris* SJ24 point at host-related differences whereby BAs may not be the key inducers for BA 7-DH-ion in the murine gut.

## Results

### In vitro bile acid transformation and impact of ^13^C-CA co-induction

Two human isolates *C. scindens* ATCC 35704 and *C. scindens* VPI 12708 and one murine isolate *E. muris* DSM 28561 (SJ24) were tested for their ability to 7-DH-ate human and mouse primary BAs *in vitro*. As expected, all three strains 7-DH-ed CA but to varying extents (Figure 2). *C. scindens* ATCC 35704 and *C. scindens* VPI 12708 showed strong 7-DH-ing activity with 97% and 80% CA conversion to 7-DH-ed BAs after 48 hours, respectively. *E. muris* SJ24, on the other hand, only converted 9% of the CA provided into 7-DH-ed forms (Figure 2). *C. scindens* ATCC 35704 produced up to 52.26 μM DCA after 32 hours with some of the DCA subsequently oxidised to 12-oxolithocholic acid (12-oxoLCA) (Figure 2A). In contrast, *C. scindens* VPI 12708 produced the highest amount of DCA after 48 hours (71.42 μM), with little to no oxidised DCA forms (0.19 μM of 3-oxoDCA at 48 hours) (Figure 2B). The lack of 12-oxo forms from the *C. scindens* VPI 12708 strain was expected since the 12α-hydroxysteroid dehydrogenase (12α-HSDH) required for this process was not detected by PCR in this strain (the full genome is currently unavailable) (data not shown). Finally, *E. muris* SJ24 only produced 8.1 μM of DCA after 48 hours with very low amounts of oxidised forms of DCA (0.21 μM of 12-oxoLCA) (Figure 2C). It is important to highlight that *E. muris* does not possess a 3α-HSDH encoded by *baiA2* which was recently identified as an important component of the CA 7-DH-ion pathway^25^. BaiA1/3 has lower affinity to CA than BaiA2^34^ and is present outside the *bai* operon^35^. Finally, all three strains also showed a modicum of 7-oxidation activity (Figure 2), resulting in the formation of 7-oxoDCA, which cannot be 7-DH-ed. Moreover, parallel experiments were performed by amending the cultures with ^13^C-CA in addition to other individual BAs in order to test whether the 7-DH-ion of CDCA, αMCA, βMCA and UDCA could be induced by CA (Supplementary Table 1). Similar results to above were obtained for the control (^13C^-CA only) experiments (Supplementary Figure 1).

**Figure 2.**
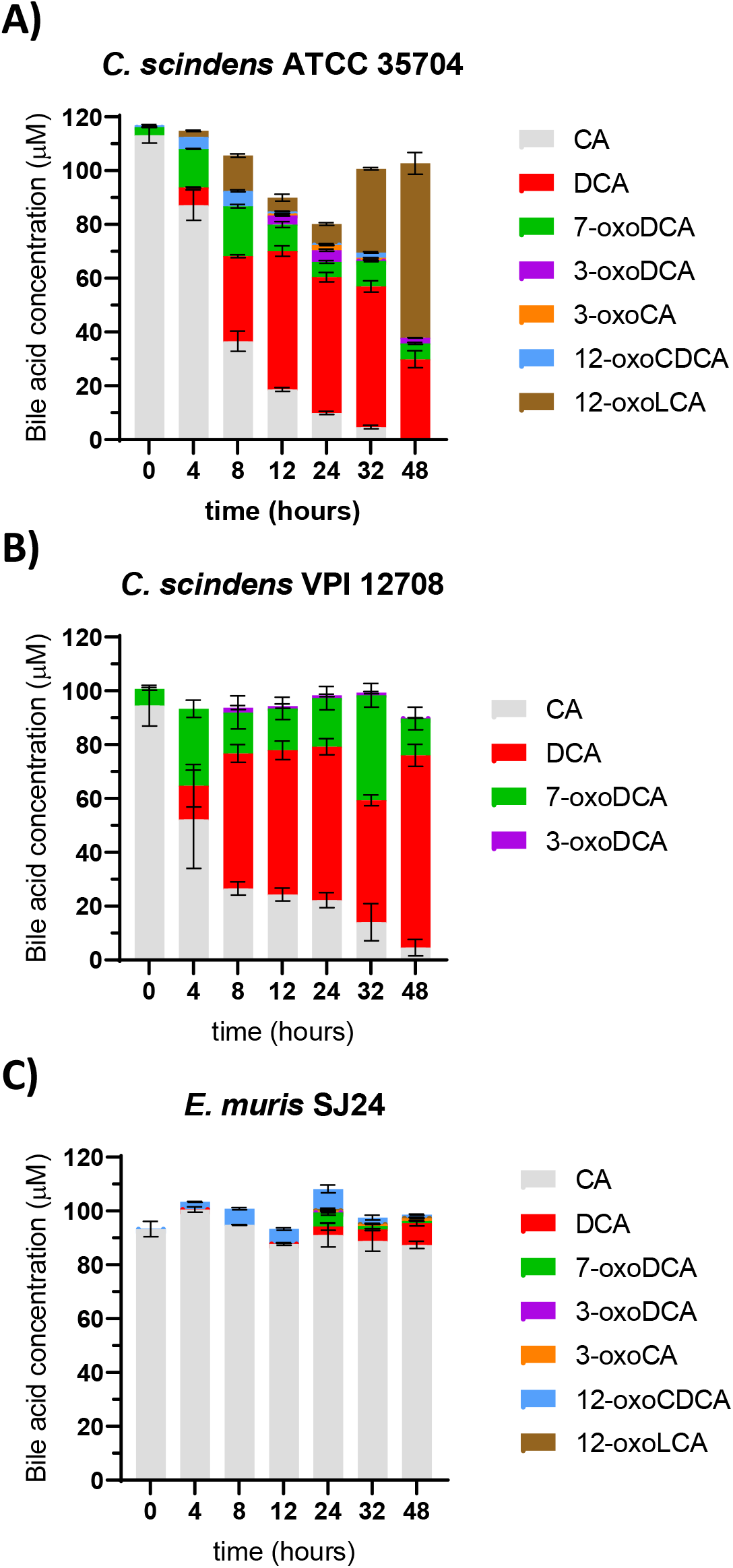
*In vitro* 7-dehydroxylation of cholic acid. The 7-dehydroxylation of CA was tested in (A) *Clostridium scindens* ATCC 35704, (B) *C. scindens* VPI 12708 and (C) *Extibacter muris* DSM 28561 (SJ24) over time. All strains were grown anaerobically in BHIS-S containing 100 μM CA. Bile acids were extracted from the suspended biomass. Error bars represent the standard deviation of the mean of biological triplicates.

CDCA 7α-dehydroxylation is very limited for all three strains. Indeed, *C. scindens* ATCC 35704 only produced 1.59 μM LCA and *C. scindens* VPI 12708 only 1.55 μM LCA (Figure 3AB), whereas for *E. muris*, no LCA was detected. The latter is in line with previous reports^21^. The amendment of ^13^C-CA significantly increased the transformation of CDCA for both *C. scindens* strains (*p-*value < 0.001 two-way ANOVA) but had no impact on *E. muris* SJ24 (Figure 3C). Indeed, the LCA yield increased to 9.77 μM for strain ATCC 35704 and to 40.4 μM for strain VPI 12708 (Figure 3AB). No change was observed for *E. muris* SJ24.

**Figure 3.**
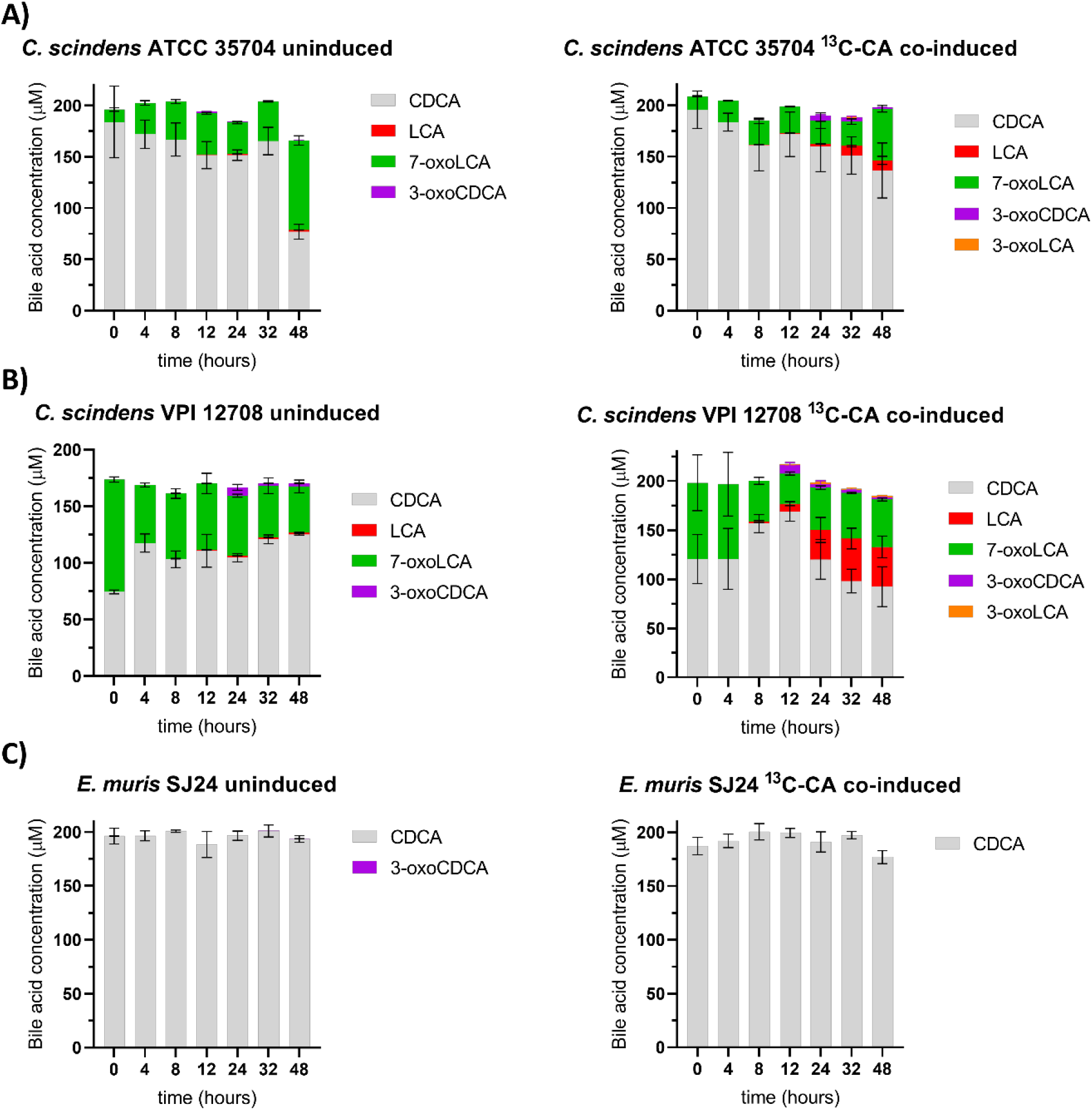
*In vitro* 7α-dehydroxylation of CDCA. The transformation of CDCA into secondary bile acids was tested with and without co-induction with 100 μM of ^13^C-CA. 200 μM of CDCA were used based on previous experiments by Marion e*t al*. (A) *Clostridium scindens* ATCC 35704, (B) *C. scindens* VPI 12708 and (C) *E. muris* DSM 28561 (SJ24). Bile acids were extracted from the suspended biomass. Error bars represent the standard deviation of the mean of biological triplicates.

As for UDCA, 7α-dehydroxylation to LCA was observed only upon amendment with ^13^C-CA (Figure 4). None of the strains exhibited any detectable level of activity from cultures that included only UDCA. 14.92 μM of LCA as well as extremely low amounts of 3-oxoLCA (with a maximum of 0.12 μM at 32 hours) were detected when *C. scindens* ATCC 35704 was co-induced. An unknown oxidised form labelled X-oxoUDCA was detected with a maximum concentration of 2.41 μM after 32 hours (Figure 4A). It is likely that this BA corresponds to 3-oxoUDCA (3-oxo-7β-hydroxy-5β-cholan-24-oic acid) as we can exclude 7-oxoLCA (the other product of oxidation of UDCA) (Figure 1). Another BA with the same ionised mass as UDCA was detected at a maximum concentration of 4.24 μM after 24 hours. We propose that this could be an isoform of UDCA with the hydroxyl group of the C3 carbon in the β conformation (3β,7β-dihydroxy-5β-cholan-24-oic acid). However, the identity of these compounds remains unconfirmed due to the lack of standards. Upon co-induction, the 7-DH-ing activity of *C. scindens* VPI 12708 was comparable to that of the co-induced ATCC strain, with 13.61 μM of LCA and 0.68 μM of 3-oxoLCA after 48 hours (Figure 4B). X-oxoUDCA was also detected at very small concentrations around 0.5 μM from 12 hours until the end of the experiment. The potential isoform of UDCA was detected at up to 9.92 μM at the 24-hour time point. Following the same trend observed with the other primary BAs, *E. muris* SJ24 did not show any detectable activity with UDCA even after co-induction. The chromatograms for the unknown bile acids can be found in Supplementary Figure 2.

**Figure 4.**
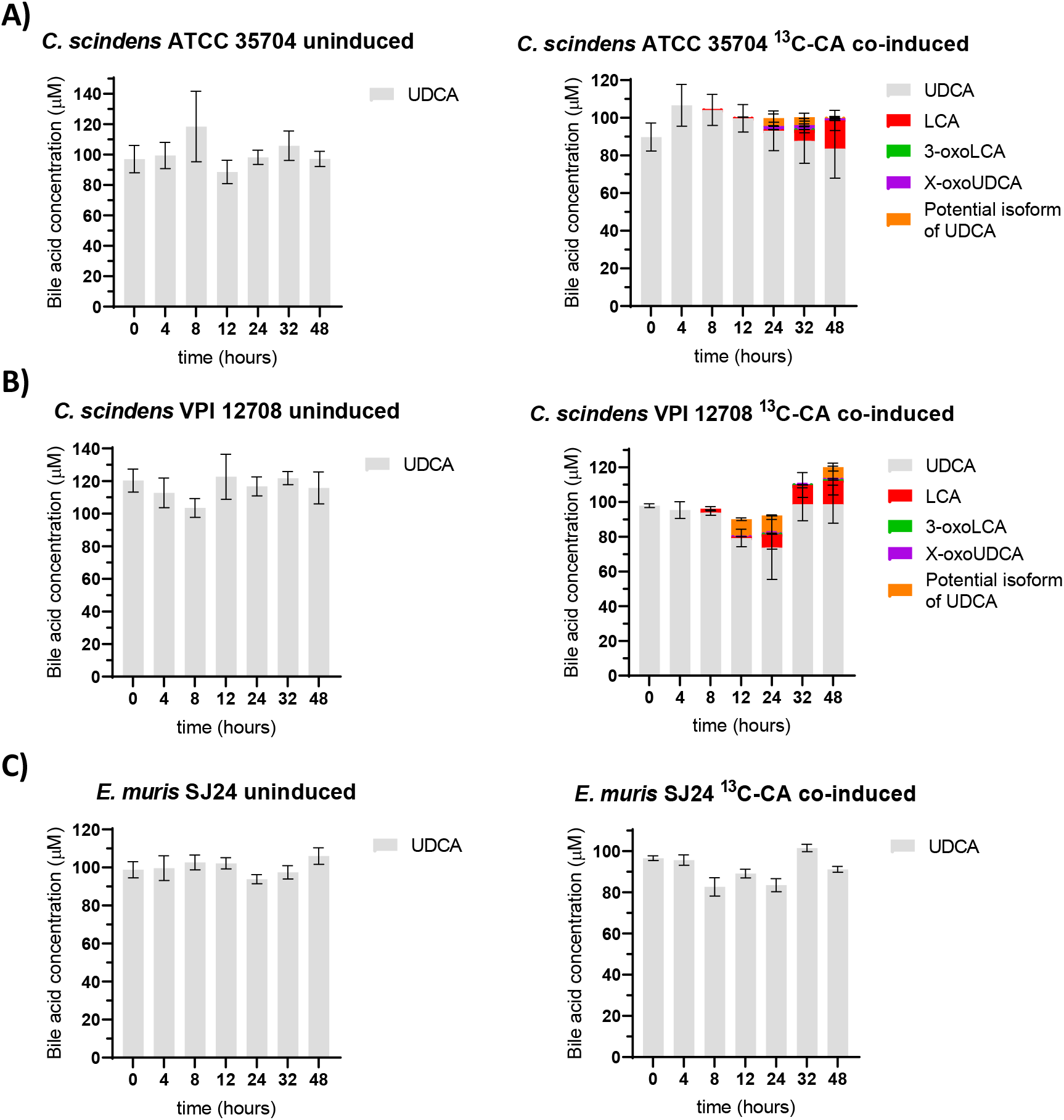
*In vitro* 7-dehydroxylation of UDCA. The transformation of 100 μM of UDCA into secondary bile acids was tested with and without co-induction with 100 μM of ^13^C-CA. (A) *Clostridium scindens* ATCC 35704, (B) *C. scindens* VPI 12708 and (C) *Extibacter muris* DSM 28561 (SJ24) were grown anaerobically in BHIS-S. Bile acids were extracted from suspended biomass. Two compounds were detected that could not be identified due to missing standards but their oxidative state can be estimated based on their ionised mass. X-oxoUDCA had the same mass as other bile acids with one ketone group and one hydroxyl group. The other unidentified compound had the same mass as UDCA and therefore it is likely to be an isoform with a 3β conformation. The retention times for these compounds was unique and therefore could not be identified further. Concentration values of the unknown BAs could only be estimated for the same reason. Error bars represent the standard deviation of the mean of biological triplicates.

As expected, neither *C. scindens* strain nor *E. muris* were capable of αMCA 7α-dehydroxylation in the absence of ^13^C-CA (Figure 5). *C. scindens* ATCC 35704 only produced minute amounts of an unknown oxo form of αMCA (labelled Y-oxoαMCA) (0.85 μM at 32 hours). Once co-induced with ^13^C-CA, 6-oxoMDCA was detected at 2.7 μM after 48 hours in the *C. scindens* ATCC 35704 culture (Figure 5A). This secondary bile acid has been 7α-DH-ed but also the hydroxyl at C6 oxidised. Moreover, several intermediates for which standards are unavailable were also detected after 48 hours. These were unknown oxidised forms of αMCA (labelled X- and Y-oxoαMCA) at concentrations not exceeding 5 μM each. A third unknown BA was detected (albeit at very low concentrations, 0.41 μM at 32 hours) with the same mass as 6-oxoMDCA, suggesting that it is an MCA species with one oxidation and one dehydroxylation. This would indicate the production of another 7α-dehydroxylated form of αMCA *in vitro* (Figure 1, Figure 5A). As for the ATCC 35704 strain, *C. scindens* VPI 12708 exhibited an increase in the quantity of products from αMCA transformation in the presence of ^13^C-CA relative to its absence (Figure 5B). This includes the 7-DH-ed BA 6-oxoMDCA that reached a concentration of 8.18 μM after 48 hours and the X- and Y-αMCA forms that were detected at maximum concentrations of 4.22 μM (4 hours) and 1.69 μM (32 hours), respectively. The aforementioned αMCA-derived bile acid with one ketone group and one dehydroxylation was also detected at a maximum concentration of 3.49 μM after 48 hours (Figure 5B). Surprisingly, *E. muris* SJ24 did not exhibit any observable 7-DH-ing activity with or without ^13^C-CA. Nevertheless, a small amount of X-oxoαMCA was detected at all time points, with a stable concentration at around 2.4 μM without and 1.9 μM with ^13^C-CA (Figure 5C). The results for βMCA were very similar to those for αMCA and are discussed in further detail in the supplementary information.

**Figure 5.**
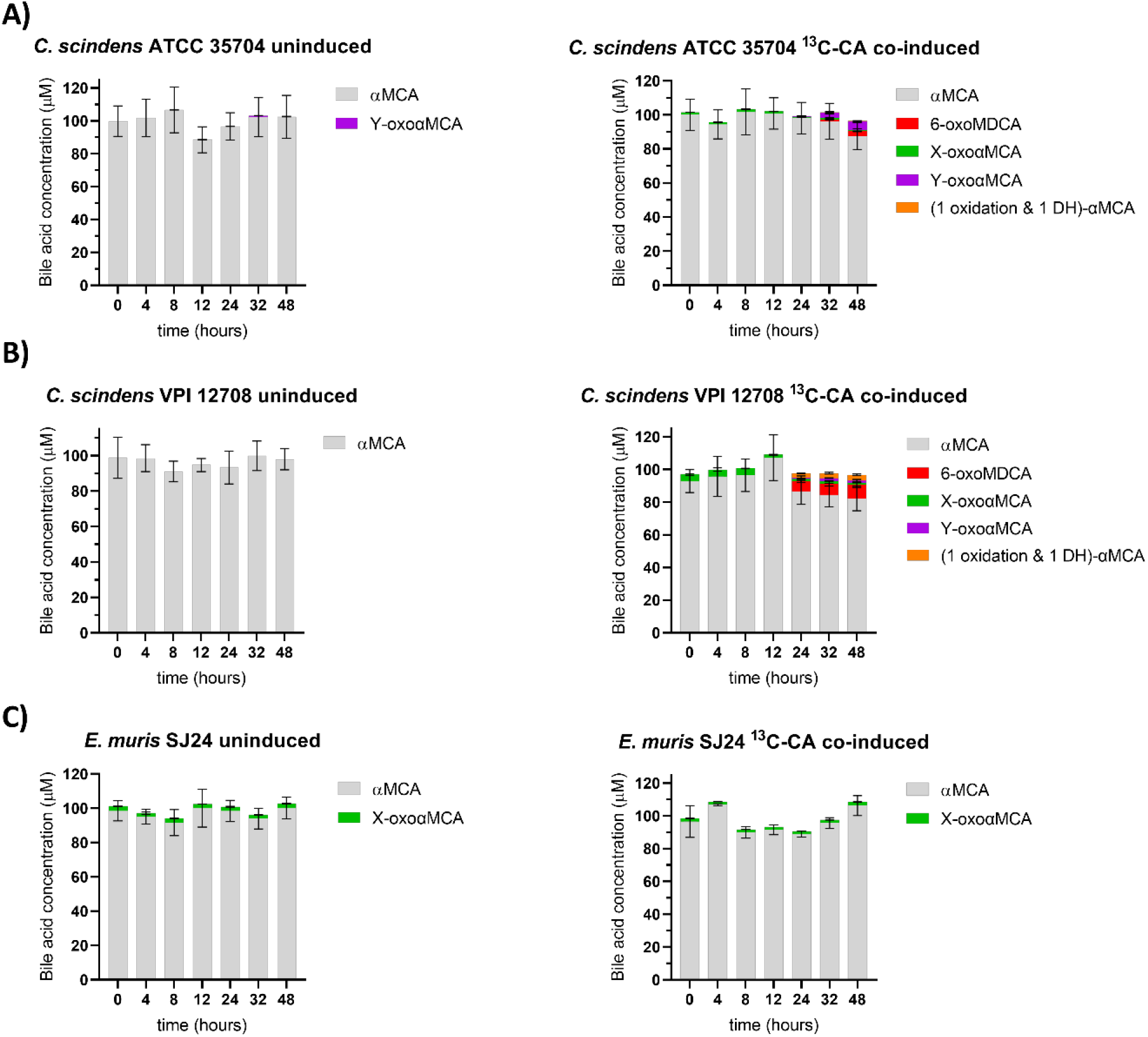
*In vitro* 7-dehydroxylation of αMCA. The transformation of 100 μM of αMCA into secondary bile acids was tested with and without co-induction with 100 μM of ^13^C-CA. (A) *Clostridium scindens* ATCC 35704, (B) *C. scindens* VPI 12708 and (C) *Extibacter muris* DSM 28561 (SJ24) were grown anaerobically in BHIS-S. Bile acids were extracted from suspended biomass. Several compounds were detected that could not be identified due to missing standards but their oxidative state can be estimated based on their ionised mass. X-or Y-oxoαMCA had the same mass as other bile acids with one ketone group and two hydroxyl groups. The other unidentified compound had the same mass as secondary bile acids that have been dehydroxylated (−1 -OH), have one ketone group and one hydroxyl group. The retention times for these compounds did not correspond to that of any known standard. Concentration values of the unknown BAs could only be semi-quantitative for the same reason. Error bars represent the standard deviation of the mean of biological triplicates.

The concentration of ^13^C-CA was also measured over time to ascertain that CA was being metabolised. It was observed to decrease until it disappeared after 48 hours in the *C. scindens* strains except in the presence of CDCA, for which the concentration decreased slowly over time. We attribute this observation to the toxicity of CDCA at that concentration^20^. On the other hand, the concentration of ^13^C-CA in *E. muris* remained stable over time and in all conditions (Supplementary Figure 3).

### Induction of bai gene expression in the presence of bile acids

In order to assess whether the amendment of ^13^C-CA to the culture induced the *bai* operon as hypothesised, the relative expression of *baiCD* and of *baiE* were measured. In addition, the expression of *baiJ* (an accessory gene) was also monitored. Gene expression was normalised using at least three reference genes and was calculated relative to the expression levels in a control group without BAs. Both *E. muris* strains, that is JM40 (DSM 28560) and SJ24 (DSM 28561), have a truncated *baiJ* gene^21^. However, while the *baiJ* pseudogene is interrupted by stop codons in strain JM40, it is not in strain SJ24, making it worthwhile to investigate *baiJ* in the latter strain (as was done in this study). Additionally, *baiO* was also analysed for *E. muris* SJ24 as an alternative accessory *bai* gene^36^.

Results show that the expression of *bai* operon genes in both *C. scindens* strains was highly upregulated as a response to exposure to CA or to CDCA but not to the other BAs (Figure 6). CA and CDCA are also the only two primary BAs for which *in vitro* 7-DH-ion data are already available^20^. Moreover, it is worth highlighting that CDCA was tested using a concentration of 200 μM vs. 100 μM for CA.

**Figure 6.**
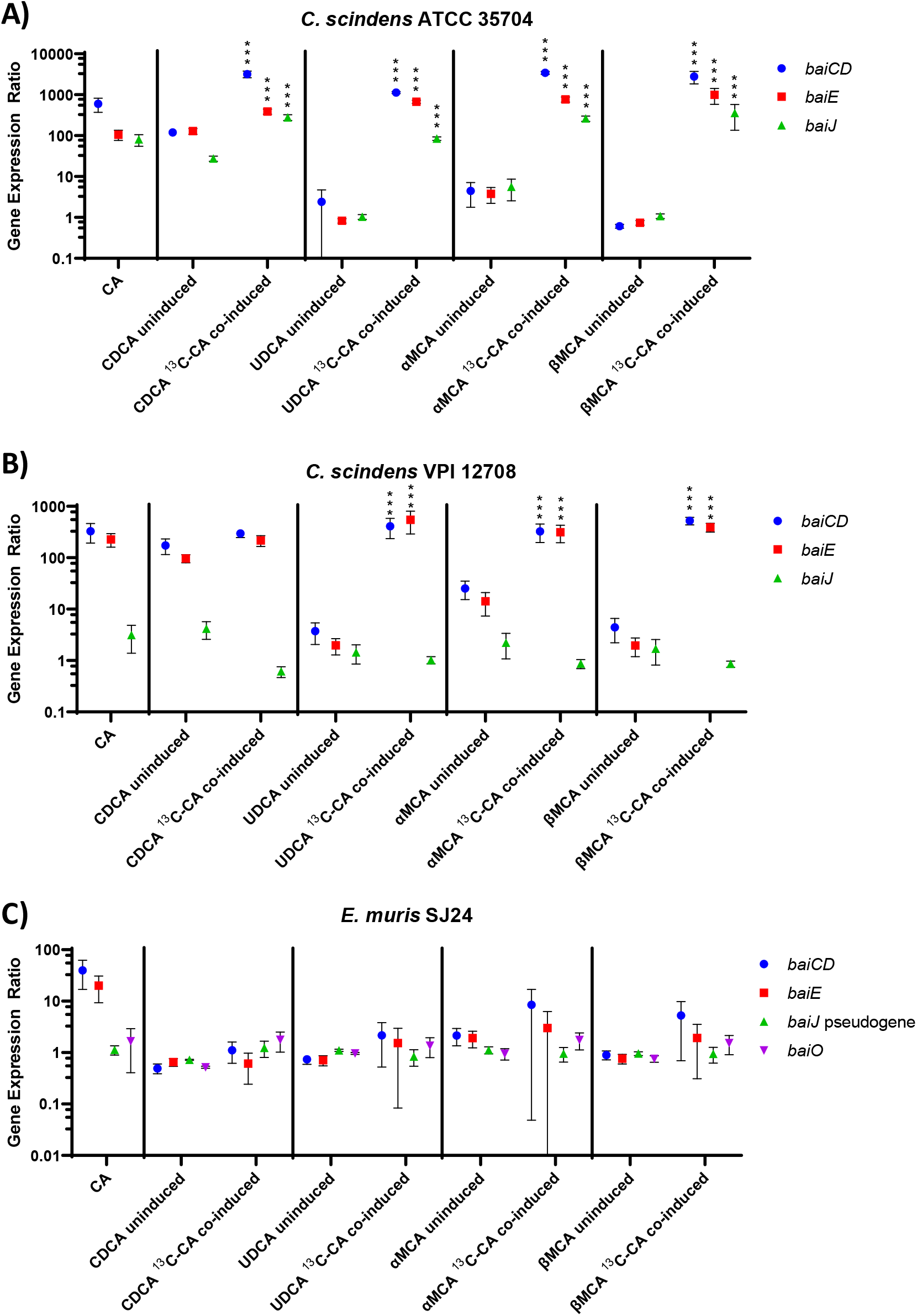
*bai* gene expression in the presence of CA (first panel) or in that of various primary BAs either uninduced or and co-induced with ^13^C-CA. Data for the CA-only condition correspond to the pooled expression results from the two sets of experiments (referred to as uninduced and co-induced) that both include this condition, as described in the text. The expression (normalised to at least three reference genes) is relative to the no BA condition including an equivalent volume of solvent (ethanol). (A) *Clostridium scindens* ATCC 35704, (B) *C. scindens* VPI 12708 and (C) *Extibacter muris* DSM 28561 (SJ24) gene expression of *baiCD, baiE* (part of the *bai* operon), and accessory genes *baiJ* and *baiO* (only in *E. muris* DSM SJ24) was measured. CA, UDCA, αMCA, βMCA and the co-inducing ^13^C-CA were used at 100 μM, CDCA was used at 200 μM. A detailed view of the *C. scindens* VPI 12708 uninduced vs 13C-CA co-induced in the CDCA group is found in Supplementary Figure 4. Coloured dots represent the average and error bars represent the standard deviation of 12 replicates. Some error bars may look elongated due to the logarithmic scale of the Y axis. (***) indicates a *p*-value < 0.001 in a linear model analysis comparing the uninduced vs ^13^C-CA co-induced factor for each *bai* gene.

For *C. scindens* ATCC 35704, the three genes tested were highly upregulated when ^13^C-CA was present along with another BA (UDCA, αMCA, or βMCA) (Figure 6A). In the CDCA dataset, statistically significant differences relative to the uninduced conditions were observed and all genes were slightly more upregulated in the presence of ^13^C-CA (*p*-value < 0.001 linear model), but *baiCD* was more so than the other genes (Figure 6A). Most interestingly, none of the other primary BAs activated the expression of *bai* genes on their own, consistent with the lack of 7-DH-ing activity with these BA substrates alone. However, the expression was brought up to levels higher than those observed for CA once co-induced (Figure 6A).

A similar pattern was observed for *C. scindens* VPI 12708 but with the significant difference being that *baiJ* was not overexpressed under any conditions (Figure 6B) (*p*-value < 0.001 linear model). In the CDCA dataset, the co-induction with ^13^C-CA had an upregulatory effect if assessed with a paired Wilcoxon test (Supplementary Figure 4) but this effect was not found to be significant when using the linear statistical model displayed in Figure 6. It is worth noting that the expression data were obtained from the mid-to late-log exponential phase (around 18 hours) when differences in activity between uninduced and co-induced conditions are not very large (Figure 3B). Similarly to *C. scindens* ATCC 35704, co-induction had a dramatic effect on the expression levels of *baiCD* and *baiE* in the presence of UDCA, αMCA, or βMCA, with upregulation reaching the expression levels observed with CA or CDCA alone (Figure 6B).

*E. muris* SJ24 showed a slight upregulation of *baiCD* and *baiE* in the CA dataset but it was not significant and did not translate to any of the other conditions (Figure 6C), consistent with its very poor 7-DH-ing activity (Figure 2). As a matter of fact, the increased *baiCD* and *baiE* gene expression ratio observed in the CA group was probably caused by one of the biological replicates which had a higher expression level than the others.

Thus, CA had a large effect on *bai* expression which was strain specific. Genes of the *bai* operon in the two *C. scindens* strains (ATCC 35704 and VPI 12708) exhibited a similar response to CA induction but the accessory *baiJ* differed in its response. It was upregulated in strain ATCC 35704 but not in strain VPI 12708. In contrast, CA had no significant effect on the expression of all the *bai* genes considered in *E. muris* SJ24.

The *rhaS*_1 gene (HDCHBGLK_01429) is immediately upstream of the *bai* operon promoter on the opposite strand and has also been proposed as bile acid-regulatory A (*barA*) due to its potential implication in *bai* regulation^7^. The expression of *rhaS1* and *rhaS2* (a copy of *rhaS1* elsewhere in the genome) was shown to have background levels across BAs (Supplementary Figure 5). This was tested in *C. scindens* ATCC 35704 without the amendment of ^13^C-CA. Results indicate that *rhaS* is not upregulated by any of the BAs tested.

Thus, the question remains about the conditions propitious for *bai* gene expression and robust 7-DH-ion in *E. muris* SJ24. We hypothesized that other mouse-specific BAs may be the key inducers

### *bai gene induction by other BAs* in *E. muris* SJ24

Because the presence of ^13^C-CA did not induce *bai* genes in *E. muris* SJ24, we tested four BA cocktails to probe whether other BAs commonly found in the BA pool could promote *bai* expression. The BA pool was divided into four cocktails: tauro-conjugated BAs, oxidised BAs, sulfonated BAs and ωMCA. The addition of these BAs did not yield the production of any detectable secondary BAs (Figure 7). A small CA concentration (<2 μM) was detected in the tauro-BA cocktail (Figure 7A) but this was likely the result of the presence of CA as an impurity in the TCA standard, as it was also detected at time 0. In the oxidised BA cocktail, 12-oxoCDCA was almost fully reduced to CA after 16 hours (Figure 7B). Small quantities of CDCA and βMCA were detected, while both are likely to be impurities from the standards used (detected at time 0), it is worth highlighting that the concentration of CDCA increased from an average of 2.74 μM (time 0) to 5.35 μM (time 24), meanwhile, the concentration of βMCA remained stable around 2 μM. No reduction of 3-oxo forms was detected, likely due to the absence of *baiA2*^25^. Finally, neither sulfonated BAs nor ωMCA were transformed by *E. muris* SJ24 in any way (Figure 7C-D). Therefore, we excluded the possibility that other BAs could induce *bai* expression and 7-DH-ing activity in *E. muris* SJ24.

**Figure 7.**
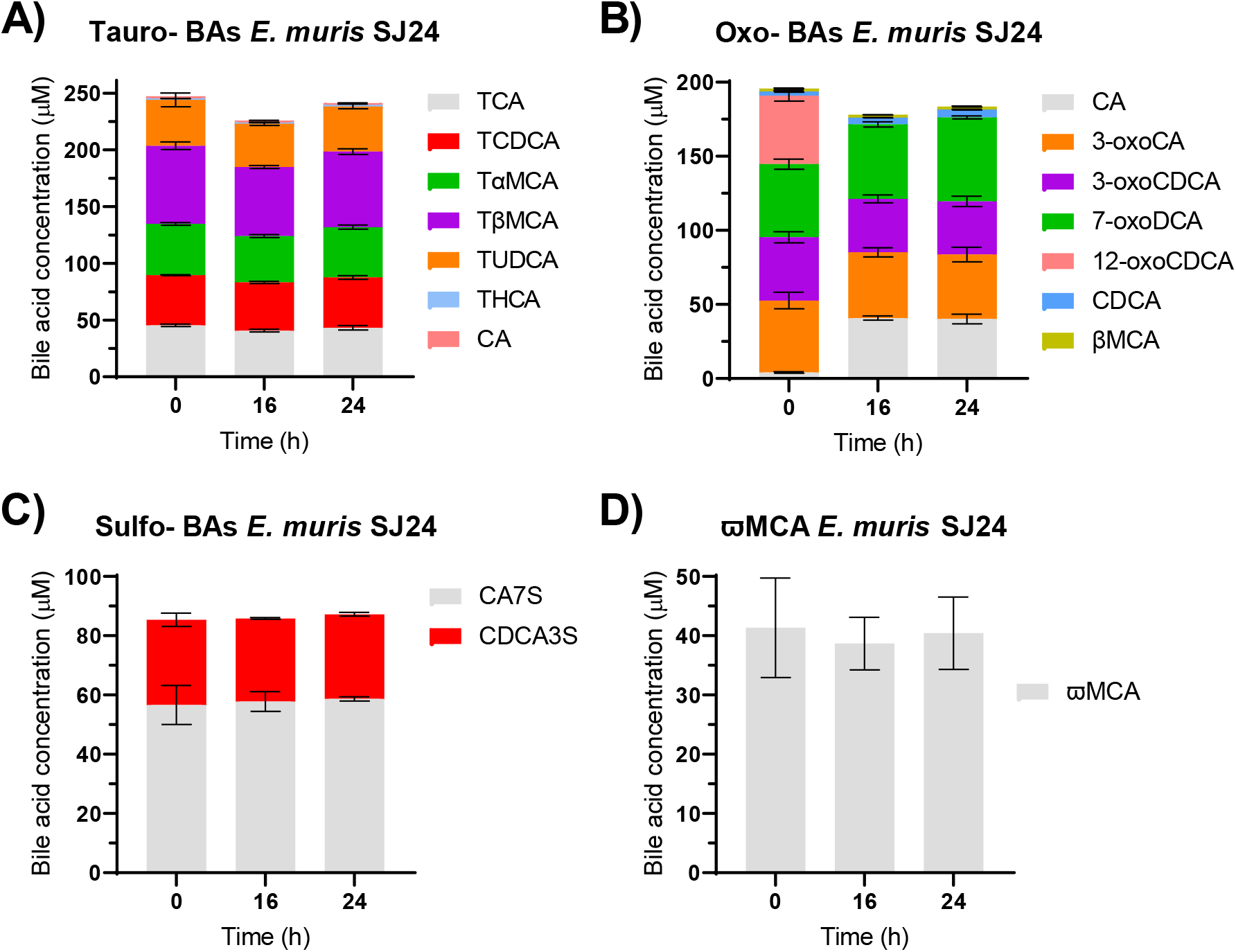
*In vitro* transformation of BA cocktails by *E. muris* DSM SJ24. The transformation of several BAs at 50 μM by *E. muris* DSM 28561 (SJ24) was tested anaerobically in 25 mL of BHIS-S. Different cocktails were prepared based on their similarity. Tauro-conjugated BAs (A). Oxidised BAs (B), Sulfonated BAs (C) or the secondary BA ωMCA (D) which is present in the murine BA pool. Three time points were taken for BA extraction from suspended biomass. An additional sample of 1 mL was taken at 16 hours for RNA extraction and RT-qPCR analysis. Minute concentrations (<2 μM) of CA (A) and βMCA (B) were detected, most likely as as impurities from the standards used as they were detected from time 0 and concentrations remained stable. Error bars represent the standard deviation of the mean of biological replicates.

The expression of *baiCD, baiE*, the pseudogene *baiJ* and *baiO* was also measured in the BA cocktail experiments and compared with a CA-only reference group. Given the lack of 7-DH-ion of the BAs within the cocktails, it is not surprising that no significant upregulation was observed in any of the BA cocktail groups when compared to the CA control. (Supplementary Figure 6A).

### Bile acid 7-DH-ion by E. muris SJ24 in the presence of mouse cecal content

CA 7-DH-ion by *E. muris* SJ24 was investigated in the presence of cecal content from either germ-free mice or a stable gnotobiotic murine model, Oligo-Mouse-Microbiota (Oligo-MM12)^37^ in order to further investigate potential non-BA triggers for 7-DH-ion. A significant fraction of CA was conjugated with Co-enzyme A (CoA) and therefore could not be detected, as there are no standards for CoA-forms. In the controls (no cecal content), the DCA concentration averaged 4.26 μM after 48 hours which corresponded to the transformation of approximately 7% of the initial CA (Figure 8A). The amendment of cecal content from germ-free mice increased the DCA produced to 7.6 μM which corresponded to 12% of the initial CA (Figure 8B). Finally, the addition of cecal content from Oligo-MM12 mice produced only 0.67 μM of DCA but 18.6 μM of 7-oxoDCA (Figure 8C). In all conditions, DCA was detected after 12 hours of incubation and gradually increased. 12-oxoCDCA was detected in all conditions, while 3-oxoCA was only found in the CA control and germ-free groups (Figure 8). The control groups of cecal content without *E. muris* SJ24 showed no change in CA concentration other than the potential conjugation with Co-A by the Oligo-MM12 mouse case (Supplementary Figure 7).

**Figure 8.**
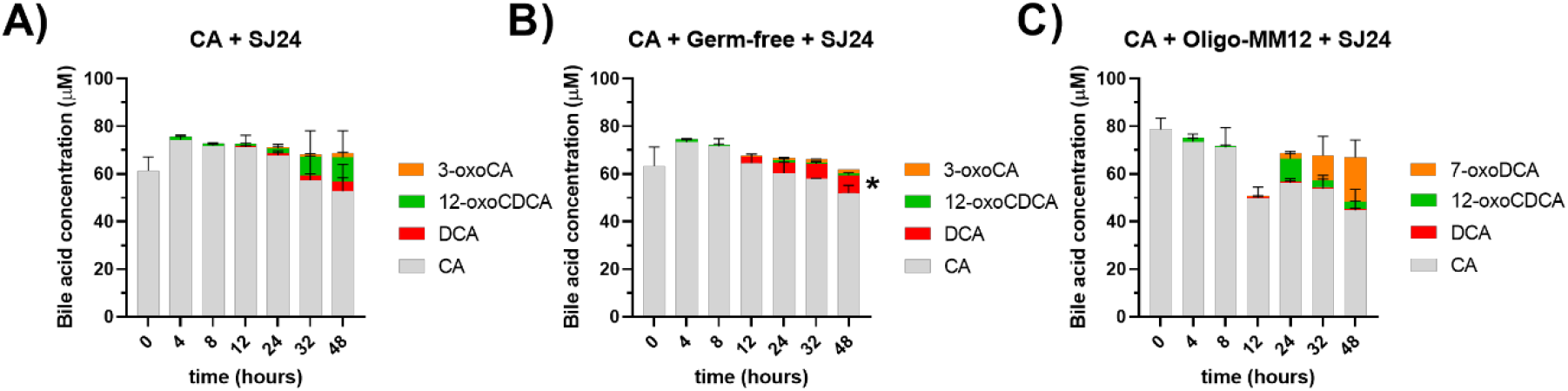
*In vitro* 7-DH-ion of CA by *E. muris* SJ24 amended with cecal content. The transformation of 100 μM of CA into secondary BAs was tested with and without the amendment of cecal content. A control group with only CA (A) was used as intra-assay reference vs (B) cecal content from germ-free mice and (C) cecal content from sDMDMm2 mice. 25 mg of cecal content were added to 25 mL of BHIS-S. The mass imbalance between the added CA (100 μM) and the measured (aprox. 80 μM) can be attributed to the presence of CoA-forms that cannot currently be quantified due to lack of standards. Error bars represent the standard deviation of biological triplicates. Data from the control groups of this experiment can be found in Supplementary Figure 7.

Despite the measurable impact on 7-DH-ion by the addition of germ-free mouse cecal content of CA 7-DH-ion, it was not sufficient to significantly upregulate *bai* expression when compared to the CA-only reference group (Supplementary Figure 6B). In both assays, the gene expression ratio of *bai* genes was never above 3.

### *E. muris* SJ24 *In vivo* 7-DH-ion and *bai* gene expression

The ability of *E. muris* strain DSM 28560 (JM40) to 7-DH-ate *in vivo* has been previously documented^21^. Here, colonisation of Oligo-MM12 mice was performed with the DSM 28561 strain (*E. muris* SJ24) to confirm 7-DH-ion *in vivo* and quantify *bai* gene expression. The bile acid composition confirms active 7-DH-ion *in vivo* in Oligo-MM12 mice. Indeed, DCA, LCA and MDCA, were exclusively identified in the sDMDMm2 + *E. muris* SJ24 group (Supplementary Figure 8).

As for the aforementioned expression assays, expression of *baiCD, baiE*, the pseudogene *baiJ* and *baiO* was quantified, normalised against at least three reference genes and calculated relative to the background signal detected from non-specific amplification in the Oligo-MM12 mice (without *E. muris* SJ24).

Results show the expression of *E*.*muris* SJ24 *bai* genes *in vivo* (Figure 9). Indeed, *baiE* was expressed at significant levels above background (gene expression ratio of 35.9) while *baiCD* and the *baiJ* pseudogene had low gene expression ratios of 1.78 and 1.38, respectively. Finally, the expression ratio of the auxiliary oxidoreductase *baiO* was situated at 4.04.

**Figure 9.**
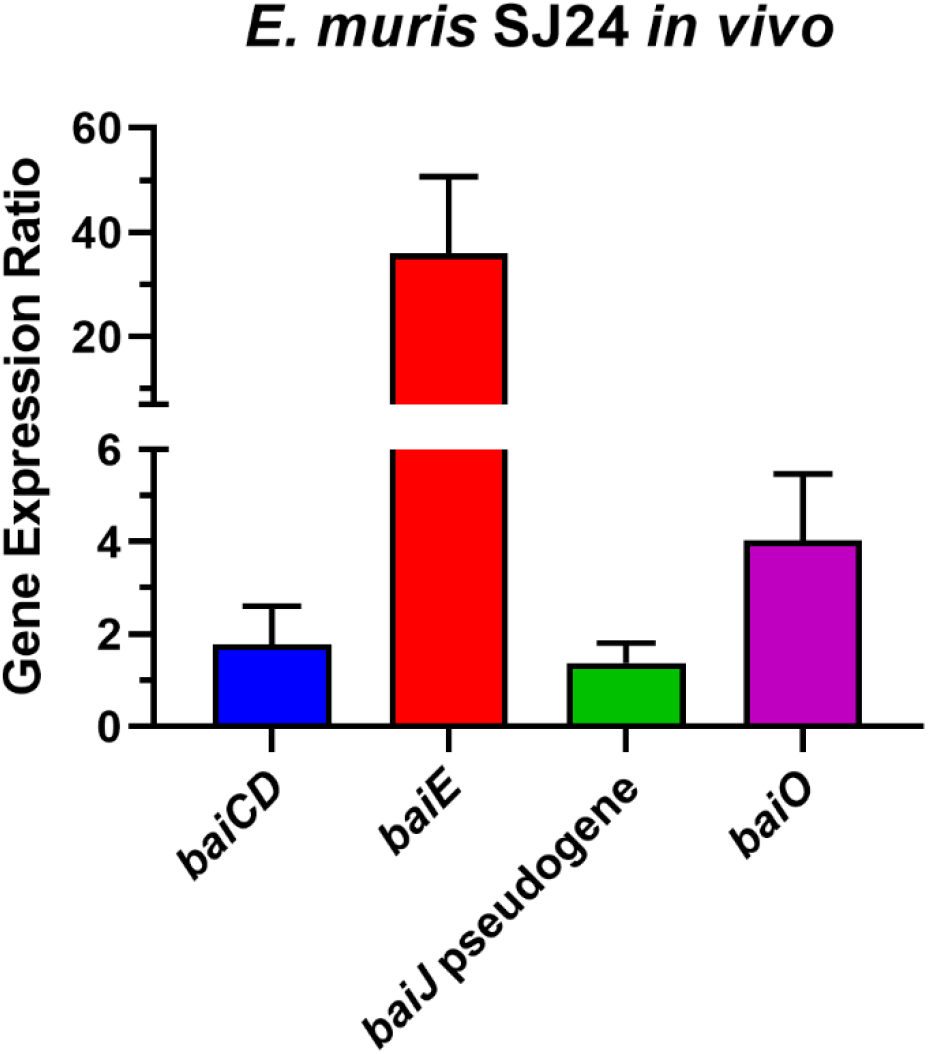
*In vivo E. muris* SJ24 *bai* expression. The expression (normalised to at least three reference genes) is relative to a non-colonised sDMDMm2 control group. Each bar represents the average value from five biological replicates with four technical replicates each. Error bars represent the standard deviation of the mean.

## Discussion

Bile acid chemistry is a relevant field in human and veterinary medicine not only because of BA detergent function during digestion but also for the wide range of roles related to host physiology and homeostasis^38,39^.

The *bai* operon was originally described in *C. scindens* VPI 12708 in 1990 and this strain has become the reference for biochemical studies of the 7-DH-ion pathway^22,35,40^. The operon (*baiBCDEFGHI*) encodes one CoA ligase (*baiB*), two oxidoreductases (*baiCD* and *baiH*), a 7-dehydratase (*baiE*), a CoA transferase (*baiF*), a transporter (*baiG*), and a putative ketosteroid isomerase (*baiI*), but not all genes are required for 7-DH-ion^25^. Expectedly from such a complex operon, comparative genomics have highlighted significant differences in the *bai* operon amongst 7-DH-ing strains^35,41^. Furthermore, a novel genetic synteny has been recently described and proven to be capable of 7-DH-ion. This novel synteny does not follow the traditional *bai* operon structure and instead breaks it down to smaller operons that cluster together^41,42^. Moreover, accessory *bai* genes have also been described across multiple strains^35,40^. These accessory genes often cluster in two operons *baiJKL* and *baiNO* which is not always complete; *C. scindens* ATCC 35704 only has *baiJ* (urocanate reductase)^32^ while the VPI 12708 strain has the full *baiJKL* set^35^. From the genome analysis, *E. muris* SJ24 has fragments of *baiJKL* as pseudogenes, while its sister strain JM40 has similar pseudogenes but interrupted by stop codons. The exact role of these genes has yet to be defined.

Out of all primary BAs, only CA has been consistently reported to be 7-DH-ed *in vitro*. In contrast, CDCA is poorly 7-DH-ed and other murine primary BAs (αMCA, βMCA or UDCA) are not at all^20,21,43–46^. We confirm that the two *C. scindens* human isolates upregulated the expression of *bai* operon genes in response to CA (Figure 6), except for *baiJ*. Indeed, we observed differences in the response of *baiJ* to CA in between the two strains. This accessory gene is annotated as an urocanate reductase and a member of the oxidoreductase family. The current CA 7-DH-ion pathway would suggest that this gene is not involved in that process in strain VPI 12708^25^, which is consistent with the lack of expression observed here (Figure 6B). Conversely, its high level of upregulation in the strain ATCC 35704 matches previously published data^32^ and suggests that *baiJ* may play a role in the 7-DH-ion of CA or other BAs in that strain. Finally, *E. muris* SJ24, the murine isolate, barely registered any *bai* gene expression response to CA or any other BA tested.

CDCA co-induction with ^13^C-CA was particularly effective in both *C. scindens* strains. The significant *bai* upregulation (Figure 6) was reflected in an increase on the 7-DH-ed products from 1 to 5% in *C. scindens* ATCC 35704 and a dramatic increase from 1 to 23% in *C. scindens* VPI 12708 (Figure 3). These data suggest that the 7-DH-ion pathway of CDCA uses the same *bai* machinery as CA and its regulation is dependent on the CA pathway. Furthermore, strain ATCC 35704 shows lower 7-DH-ion of CDCA (Figure 3) and less effective 7-DH-ion of CA (Figure 2 and Supplementary Figure 1) as compared to strain VPI 12708 despite a significant upregulation in *baiJ* expression.

UDCA is the 7β isomer of CDCA and is reported to have significant therapeutic properties^2,7,47^. In humans, UDCA is synthesised from CDCA by gut microbes containing 7β-HSDHs^7,26,48,49^ and is thus, a secondary BA; in mice, it is produced by the liver, thus, it is a primary BA, and is used as a precursor for βMCA^15,16^. Nevertheless, colonisation of mice lacking 7-dehydroxylating bacteria with 7-DH-ing bacteria increases UDCA levels, implying that the gut microbiome also plays a significant role in UDCA production by mice^14^. Despite the importance of this BA for medical applications, little is known about the capacity of bacteria to 7β-dehydroxylate UDCA *in vitro*. Previous work has reported no transformation by *C. scindens* strain ATCC 35704^20^. In accordance with our hypothesis, co-induction with ^13^C-CA not only greatly upregulated *bai* gene expression (Figure 6B) but also provided evidence of UDCA 7β-dehydroxylation as LCA was detected in the culture (Figure 4). Indeed, the amount of 7-DH-ed products upon co-induction increased from 0 to 15 and 16% in *C. scindens* strains ATCC 35704 and VPI 12708, respectively. The absence of CDCA in the culture suggests that UDCA was not epimerised to CDCA and subsequently 7α-dehydroxylated. Moreover, two unidentified compounds consistent with 7β-dehydroxylation were detected, providing further evidence of the 7β-dehydroxylation of UDCA and its associated unique set of intermediates. The oxidised intermediate is very likely to be 3-oxoUDCA (3-oxo-7β-hydroxy-5β-cholan-24-oic acid), as based on the mass, the only alternative would have been 7-oxoLCA, which can be excluded as it is one the standards within our collection (Figure 4). The other unknown compound had the same mass as UDCA albeit a different retention time, suggesting this could be an iso-form of UDCA. It follows that this would be 3β-UDCA, tentatively named isoUDCA (3β,7β-dihydroxy-5β-cholan-24-oic acid). Isoforms are well known in the BA pool and particularly in CDCA-related intermediates^20,25,33,50^ so it is likely that UDCA follows a similar pattern. Despite the production of unknown intermediates, the activity response to co-induction suggests that UDCA 7β-dehydroxylation uses the same Bai machinery as CDCA 7-DH-ion.

7-DH-ion activity was uncovered for αMCA and βMCA, for the first time and it yielded several intermediate BAs that we were not able to fully characterise due to the lack of appropriate standards.

All human and mouse BAs share a backbone of four rings (Figure 1), this makes mass fractionation in the mass spectrometer unsuitable for identification. Therefore, we currently rely on comparison of ionised mass and retention time to standards. However, several assumptions can be made to speculate what these compounds could be. MCAs could be oxidised at the C-3, C-6 or C-7 position (Figure 1). A C-7 oxidation would yield 7-oxoMDCA regardless of the primary MCA. Meanwhile, other oxidations could be differentiated by the α or β conformation of the C-7 hydroxyl. It is possible that one of the intermediates that we detected was 7-oxoMDCA but none shared retention times across MCAs, meaning that different intermediates were produced for each of the two MCA substrates (Supplementary Figure 2). Considering that distinct oxidised intermediates were detected for the two MCAs, we hypothesise that those are the 3-oxo and 6-oxo forms of αMCA and βMCA (3-oxoαMCA: 3-oxo-6β,7α-dihydroxy-5β-cholan-24-oic acid; 6-oxoαMCA: 6-oxo-3α,7α-dihydroxy-5β-cholan-24-oic acid; 3-oxoβMCA: 3-oxo-6β,7β-dihydroxy-5β-cholan-24-oic acid; and 6-oxoβMCA: 6-oxo-3α,7β-dihydroxy-5β-cholan-24-oic acid). A third intermediate was also detected from both MCAs, with the ionised mass corresponding to secondary BAs with one dehydroxylation and a ketone group (e.g., 7-oxoLCA) (Figure 5 & Supplementary Figure 9). Three options are plausible: 1) A dehydroxylation at the C-3 position. This would yield a novel family of BAs with a 6β- and 7α/β-hydroxyls which is highly unlikely as it would have been identified previously by the multiple studies investigating the murine BA pool^15,16,51–54^. 2) An oxidation paired with a 6-dehydroxylation would yield 3-oxoCDCA, 7-oxoLCA, or 3-oxoUDCA. 3-oxoCDCA and 7-oxoLCA were included as standards in our analysis (Supplementary Table 3) and would have been detected if present. Meanwhile, the retention time of this compound is distinct from that of the compound proposed to be 3-oxoUDCA from the transformation of UDCA (see above and Supplementary Figure 2). Thus, this is not likely to be 3-oxoUDCA. 3) A 7-dehydroxylation could allow for ketone groups at the C-3 and C-6 positions. 6-oxoMDCA was available as a standard but, the second option, 3-oxoMDCA (3-oxo-6β-hydroxy-5β-cholan-24-oic acid), was not. It is therefore possible that this compound corresponds to 3-oxoMDCA, but this remains to be confirmed.

The mouse BA pool is significantly more diverse than that of humans due to the primary production of muricholic acids (with a hydroxyl group at the C-6 position) and of UDCA. The murine secondary BA pool includes DCA and LCA but also MDCA and its 6α counterpart, hyodeoxycholic acid (HDCA). Furthermore, mice can rehydroxylate TDCA back into TCA in the liver^15^ which magnifies the differences between mouse and human BA pools. In general, the secondary BAs derived from muricholic acids seem to be in low abundance in the gut, hinting at the difficulty in 7-DH-ing these BAs. This is perhaps the reason why primary BAs such as βMCA are highly abundant in the mouse BA pool^55^. While we initially hypothesised that the lack of 7-DH-ion activity for αMCA and βMCA was due to the lack of *bai* gene expression, our co-induced data show that even *bai* gene expression in the *C. scindens* strains is insufficient for the production of MDCA^21^, and results in the detection of potential oxidized versions of that BA (Figure 5 and Supplementary Figure 9).

As reported above, the murine strain SJ24 has shown no significant *bai* upregulation nor 7-DH-ion *in vitro* either in the presence or absence of ^13^C-CA. To investigate the underlying reasons for this lack of activity, we considered three additional conditions: (a) various BA mixtures, to determine whether *bai* gene expression was controlled by another (or several other) murine BAs; (b) *in vitro* in the presence of Germ-free or Oligo-MM12 cecal content to ascertain whether the presence of other gut bacteria or signalling molecules from the host itself induced 7-DH-ion; or (c) in the Oligo-MM12 environment, to confirm the activity of strain SJ24 *in vivo*.

First, the *in vivo* condition exhibited high upregulation of *baiE* relative to the background, non-colonised control (Figure 9) and 7-DH-ed BAs were detected in the BA pool from the same samples (Supplementary Figure 8). On the other hand, *baiCD* did not show evidence of upregulation. It is possible that the expression of *baiCD* by *E. muris* is constitutive while *baiE*, a 7α-dehydratase^56^, presents a higher expression ratio, perhaps due to the smaller size of the protein (169 aa in *E. muris* SJ24), intra-operon regulatory elements, or differential mRNA half-life^57^. The expression of *baiO*, a different oxidoreductase, was slightly above that of *baiCD*. Whilst the gene expression ratios remained low in both instances, a role in 7-DH-ion by this protein cannot be ruled out. Thus, *E. muris* SJ24 is capable of *in vivo* 7-DH-ion, although its *bai* machinery may require additional genes that have not been identified in this study. However, the exact trigger for *in vivo* levels of 7-DH-ion from *E. muris* still remained elusive at this point.

To elucidate that question, we tested all the BAs detected in the Oligo-MM12 environment, and found no evidence of 7-DH-ing activity (Figure 7), excluding the possibility that non-CA BA triggered *bai* gene expression in *E. muris*.

However, when SJ24 was grown in the presence of cecal content from germ-free mice, its 7-DH-ing activity increased (Figure 8) despite *bai* gene expression not increasing significantly (Supplementary Figure 6B). Indeed, the amendment of cecal content from germ-free mice (Figure 8B) resulted in a significant increase from 7 to 13% of 7-DH-ed products, coupled with a decrease of the abundance of the 12-oxoCDCA (3α,7α-dihydroxy-12-oxo-5β-cholanic acid) intermediate. Interestingly, the co-cultivation of *E. muris* with the non-sterile cecal-content from Oligo-MM12 mice produced high amounts of 7-oxoDCA (Figure 8C) while CA was not transformed by the same cecal content in the absence of strain SJ24 (Supplementary Figure 7). This could suggest an interaction between *E. muris* and the gnotobiotic community in which the former would promote the 7-oxidation of CA by the latter, known to harbor 7-HSDHs^58^. This interaction appears to be exclusive to the *in vitro* environment since the *in vivo* BA data show lower concentrations of 7-oxoDCA than DCA (Supplementary Figure 8).

The evidence presented in this study shows that the role of the host, presumably through signalling, is a critical element for effective 7-DH-ion by *E. muris* and the regulatory mechanisms of this secondary BA transformation is dramatically different amongst bacterial species. The results also highlight the significant differences in 7-DH-ion between human and mouse isolates but also between the *in vitro* and *in vivo* environments.

To further highlight the differences, the human isolates showed marginal activity for αMCA and βMCA upon co-induction. 7-DH-ed forms such as 6-oxoMDCA were detected (Figure 5 & Supplementary Figure 9) but the full 7-dehydroxylation to MDCA was not observed. These data add further evidence that other elements besides the *bai* operon (and *baiJ*) might be needed to completely 7-dehydroxylate MCAs. Perhaps, a missing 6β-HSDH gene would be required for complete 7-DH-ion. The apparent simplicity of 7-DH-ion regulation from human isolates compared to that of *E. muris* could be due to the more diverse diet of humans than mice. It has been observed that a less diverse diet can overstimulate the BA pool in humans and increase the incidence of colorectal cancer^59^. The natural mouse diet is less diverse than that of humans and their initial lactation period (a monotrophic diet) plays a much stronger role in the mouse lifespan^60^. In these circumstances, a strong regulation of 7-DH-ion might be an important mechanism to prevent BA pool unbalances. Nevertheless, much more data on the 7-DH-ion mechanisms of various strains with particular focus on isolates from the mouse and other animal models is required to investigate the potential differences in 7-DH-ion regulation.

## Conclusion

The findings presented here are fourfold. First, we demonstrated that the previously reported^32,33^ strong upregulation of *bai* genes by CA increases the extent of 7-DH-ion of other primary BAs. This was particularly true for UDCA which has been reported to be converted to LCA *in vitro* for the first time, with important differences in the 7-DH-ing capabilities amongst strains.

Secondly, the upregulation of *bai* genes exhibited strain-specific differences. *C. scindens* ATCC 35704 upregulated the *bai* operon genes *baiCD* and *baiE* as well as the *bai* accessory gene *baiJ*. While for *C. scindens* VPI 12708, it upregulated the *bai* operon genes, but *baiJ* expression was found to be at a background level. This is consistent with the lack of involvement of *baiJ* in 7-DH-ion in strain VPI 12708^25^. *E. muris* SJ24 was the third bacterium tested, a murine isolate with *in vivo* 7-DH-ing capabilities (Supplementary Figure 8). Strain SJ24 showed weak *in vitro* 7-DH-ion of CA and no upregulation of any of the *bai* genes tested. The activity of this strain was promoted by the addition of germ-free cecal content but not cecal content from Oligo-MM12 colonized mice. This result suggests that host signalling is required for efficient 7-DH-ing activity by strain SJ24 but that the presence of a minimum microbiome (12 strains) inhibits this activity potentially due to the promotion of 7-HSDH activity from the microbiome *in vitro*. Unravelling the controls on BA 7-DH-ion by *E. muris* requires further investigation.

Thirdly, *C. scindens* human isolates can partially 7-dehydroxylate MCAs, leading to the formation of oxidized MDCA at the C-6 position. Therefore, an enzyme capable of reducing this compound, e.g., a 6β-HSDH, is required to achieve the end point of 7-DH-ion that is observed *in vivo*, namely MDCA. To date, such protein has not been identified in any microorganism.

Finally, the mechanism of 7-DH-ion regulation differs significantly between murine- and human-derived strains, which could be due to the nature of the host. Human isolates showcase a system governed by CA, while murine isolates appear to utilise a BA-independent signal present in the lumen.

In conclusion, these data provide novel insights into the intricacies of 7-DH-ion and the significant differences amongst 7-DH-ing bacteria. The CA-dependent co-induction can be attributed to the abundance of this compound in the BA pool of humans. However, the inducing factor for *E. muris* activity remains elusive despite evidence suggesting that it is host-derived. Moreover, multiple novel BAs were observed and their identity surmised. Further work with these and other strains is required to investigate the 7-DH-ion pathway of CDCA, αMCA, βMCA and UDCA as well as to explore the strain-specific differences regarding the 7-DH-ion pathway of CA.

## Materials and Methods

### Bacterial strains and growth conditions

The strains used were *Clostridium scindens* ATCC 35704, *Clostridium scindens* VPI 12708 and *Extibacter muris* DSM 28561 (SJ24), this strain was chosen instead of *E. muris* DSM 28560 (JM40) due to its ability to grow faster *in vitro* (24h vs 48h, data not shown). Bacteria were grown in Brain Heart Infusion Supplement – Salts (BHI-S) medium, consisting of 37g BHI, 1g L-cysteine, 5g yeast extract, 2.5g fructose, 50mL salts solution (0.2g CaCl_2_, 0.2 MgSO_4_, 1g K_2_HPO_4_, 1g KH_2_PO_4_, 10g NAHCO_3_ and 2g NaCl per L of ddH_2_O) per L ddH_2_O. The salts solution and media were sterilised by autoclaving. Static growth was carried out at 37°C in an anoxic chamber (Coy Laboratory Products, 95% N_2_, 5% H_2_). A pre-inoculum was prepared from glycerol stocks (using BHIS-S) before inoculating 25 mL of BHIS-S in Falcon tubes at a starting OD_600_ of 0.05. Results of the growth curves for the experiments can be found in the Supplementary Information (Supplementary Figure 10).

### *In vitro* 7-dehydroxylation assays

Bacteria were grown in presence of BAs: CA (100 μM), CDCA (200 μM), αMCA (100 μM), βMCA (100 μM) and UDCA (100 μM) or the same volume of ethanol (solvent control). A sterile control (media with ethanol solvent) was also included. Co-induction was performed by adding an additional 100 μM of ^13^C-CA to CDCA, UDCA, αMCA, or βMCA). ^13^C-CA was chosen so its transformation products could be separated from those of the other primary BAs during the quantification process. All uninduced experiments were conducted at the same time while all the co-induced ones were run simultaneously at a different time. Both experiments included a condition consisting of the amendment of only 100 μM CA, which allows for comparison of the results from co-induced and uninduced experiments. Growth was monitored by periodically measuring the OD_600_. During the main time points (0, 4, 8, 12, 24, 32 and 48 hours), 1mL samples were collected for BA extraction in a 2 mL bead-beating resistant tube. All conditions were performed in triplicates.

Four BA cocktails were prepared for the assays with *E. muris* SJ24 based on the BAs often present in a meaningful amount within the BA pool of mice. All BAs within the cocktails were at 50 μM. The Tauro-conjugated cocktail included TCA, TCDCA, TαMCA, TβMCA, TUDCA and THCA. The Oxo-cocktail included 3-oxoCA, 3-oxoCDCA, 7-oxoDCA and 12-oxoCDCA. The Sulfo-cocktail included CA7S and CDCA3S. Finally, the last group was only comprised of 50 μM of ωMCA. All BAs were diluted in ethanol or methanol depending on their solubility. Three time points were collected (0, 16 and 24 hours).

Furthermore, *E. muris* SJ24 was also amended with cecal content to test its implications over 7-DH-ion *in vitro*. For this assay, 25 mg of freeze-dried cecal content from germ-free mice or 25 mg of frozen content with 5% glycerol (v/v) from sDMDMm2 mice were added to 25 mL of BHIS-S with 100 μM CA. The control conditions for this experiment were a group with 100 μM CA but no additional cecal content and two more groups with each respective type of cecal content but no *E. muris* SJ24. Seven time points were taken (0, 4, 8, 12, 24, 32 and 48). RNA sample collection was particularly early for this experiment (4-hour time point) due to a faster-than-usual growth (Supplementary Figure 10).

### Bile acid extraction

Samples were vacuum dried overnight (ON). Approximately 450 mg of 0.5 mm zirconium beads were added to the dried samples as well as 500 μL ice-cold alkaline acetonitrile (acetonitrile – 25% ammonia 4:1 v/v) and 100 μL of ISTD solution (CA-d_4_, CDCA-d_4_, TCA-d_4_, TUDCA-d_4_, DCA-d_4_ and LCA-d_4_, each at 100 μM in methanol). Samples were homogenised in a Precellys 24 Tissue Homogenizer (Bertin Instruments, Montigny-le-Bretonneux, France) at 6500 rpm 3x 30” beat 30” rest. Samples were vortexed for 1 hour and centrifugated for 15 minutes at 16000 rcf at room temperature. Approximately 500 μL of suspension was carefully collected over the beads level and transferred into a new 1.5 mL epi tube which was then vacuum dried overnight. Finally, the samples were reconstituted in 1 mL of ammonium acetate [5mM] – methanol (50:50 v/v) and a 1:20 dilution with the same solvent was prepared in LC-MS glass vials, ready for injection.

### RNA extraction and reverse transcription

1 mL of sample was collected in a 15 mL falcon tube during the mid-log to late-log phase for the RNA extraction. The sample was stored with RNAprotect following the manufacturer protocol (Protocol 5 from RNAprotect Bacteria Reagent Handbook 01/2020, Qiagen) at -80°C until processed. All conditions were performed in triplicates. Lysis and RNA purifications were done using the RNeasy Mini Kit (Qiagen, Hilden, Germany). Bacterial lysis was performed following Protocol 5: Enzymatic Lysis, Proteinase K Digestion and Mechanical Disruption of Bacteria (RNAprotect Bacteria Reagent Handbook 01/2020, Qiagen), with 20 μL of proteinase K for each sample and the required volumes for a number of bacteria <7.5 × 10^8^. The cell lysis was performed using a Precellys 24 Tissue Homogenizer (Bertin Instruments, Montigny-le-Bretonneux, France) at 6500 rpm 3x 10 seconds beat 10 seconds rest. RNA purification was performed following Protocol 7: Purification of Total RNA from Bacterial Lysate using the RNeasy Mini Kit. Centrifugations were carried out at 15000 rcf except for the 2 min centrifugation which was done at 18000 rcf.

Purified RNA was further subject to a DNase treatment using the RQ DNase I (Promega, Madison, WI, USA) following the manufacturer protocol with small modifications: The final volume was adjusted to a 100 μL and incubation was extended to 1 hour at 37°C. The treated RNA was cleaned-up using the RNeasy Mini Kit (Qiagen, Hilden, Germany) following the RNA Clean-up protocol from the manufacturer (RNeasy Mini Handbook 10/2019) with the 2 min centrifugation done at 18000 rcf. Concentration and purity of RNA was measured with a NanoDrop One (Thermo Fisher Scientific, Waltham, MA, USA).

100 ng of RNA was reverse transcribed into cDNA using the GoScript™ Reverse Transcription Mix, Random Primers (Promega, Madison, WI, USA) following the manufacturer protocol. The process was done in duplicates with one group using water instead of the reaction buffer as a non-reverse transcription control (NRT).

### Reverse transcription quantitative PCR (RT-qPCR)

RT-qPCRs were prepared using the Myra liquid handling system (Bio Molecular Systems, software version 1.6.26) and performed using the Magnetic induction cycler (Mic) platform (Bio Molecular Systems, Upper Coomera, QLD, Australia) with the micPCR software (v2.10.5).

The list of primers used can be found in Supplementary table 2. Samples were prepared with the SensiFAST SYBR No-ROX Kit (Meridian Bioscience, Cincinnati, OH, USA) at a final volume of 10 μL. All runs were performed with the following program, with small modifications: Initial hold at 95°C for 5 minutes with a cycle of 95°C for 5 seconds, 54.5°C for 20 seconds (54.1°C for *E. muris* SJ24) and 72°C for 9 seconds. 40 cycles were done for *C. scindens* ATCC 35704 and 50 for *C. scindens* VPI 12708 and *E. muris* SJ24. The melting curve, temperature control and acquisition settings were left as default. The quantification was done using three or more reference genes (Supplementary Table 2) based on their expression stability across conditions. NRTs as well as no template controls (NTCs) were included to check for residual DNA or contaminations. Four technical replicates were done for each biological replicate. Note that expression data presented in Figure 6 for the CA-only condition (labelled CA) correspond to the pooled expression results (for the condition in which only 100 μM Ca was added) from the two sets of experiments presented above (referred to as the co-induced and the uninduced experiments, respectively). The *in vivo* expression data presented in Figure 9 was normalised against the background signal detected from an uncolonised Oligo-MM12 control group.

### Liquid chromatography – mass spectrometry (LC-MS)

The quantitative method was performed on an Agilent ultrahigh-performance liquid chromatography 1290 series coupled in tandem to an Agilent 6530 Accurate-Mass Q-TOF mass spectrometer. The separation was done on a Zorbax Eclipse Plus C18 column (2.1 × 100mm, 1.8 μm) and a guard column Zorbax Eclipse Plus C18 (2.1 × 5mm, 1.8 μm) both provided by Agilent technologies (Santa Clara, CA, USA). The column compartment was kept heated at 50°C. Two different solutions were used as eluents: ammonium acetate [5mM] in water as mobile phase A and pure acetonitrile as mobile phase B. A constant flow of 0.4 mL/min was maintained over 26 minutes of run time with the following gradient (expressed in eluent B percentage): 0-5.5 min, constant 21.5% B; 5.5-6 min, 21.5-24.5% B; 6-10 min, 24.5-25% B; 10-10.5 min, 25-29% B; 10.5-14.5 min, isocratic 29% B; 14.5-15 min, 29-40% B; 15-18 min, 40-45% B; 18-20.5 min, 45-95% B; 20.5-23 min, constant 95% B; 23-23.1 min, 95-21.5% B;

23.10-26 min, isocratic 21.50% B. The system equilibration was implemented at the end of the gradient for 3 minutes in initial conditions. The autosampler temperature was maintained at 10°C and the injection volume was 5μL. The ionisation mode was operated in negative mode for the detection using the Dual AJS Jet stream ESI Assembly. The QTOF acquisition settings were configured in 4GHz high-resolution mode (resolution 17000 FWHM at m/z 1000), data storage in profile mode and the high-resolution full MS chromatograms were acquired over the range of m/z 100-1700 at a rate of 3 spectra/s. The mass spectrometer was calibrated in negative mode using ESI-L solution from Agilent technologies every 6 hours to maintain the best possible mass accuracy. Source parameters were setup as follows: drying gas flow, 8 L/min; gas temperature, 300°C; nebulizer pressure, 35psi; capillary voltage, 3500V; nozzle voltage, 1000V. Data were processed afterwards using the MassHunter Quantitative software and MassHunter Qualitative software to control the mass accuracy for each run. In the quantitative method, 42 bile acids were quantified by calibration curves (Supplementary Table 3). The quantification was corrected by addition of internal standards in all samples and calibration levels. Extracted ion chromatograms were generated using a retention time window of ± 1.5 min and a mass extraction window of ± 30ppm around the theoretical mass of the targeted bile acid. Unknown BAs were identified when found within the retention time window of a standard with the same ionised mass. Approximate quantification of these unknown BAs was done by using the nearest standard (by retention time) with the same ionised mass.

### Animals and Ethics Statement

sDMDMm2^61^ mice were housed in the Clean Mouse Facility (CMF, Department of Clinical Research) of the University of Bern. Animal experiments were performed in accordance with the Swiss Federal and the Bernese Cantonal regulations and were approved by the Bernese Cantonal ethical committee for animal experiments under the license number BE82/13.

### *In vivo* colonisation with *E. muris* SJ24

A cohort of nine sDMDMm2^61^ mice were used. Four mice were dedicated to an uncolonized control group and the remaining five were colonised with *E. muris* DSM 28561 (SJ24). sDMDMm2 animals to be colonised were imported from breeding isolators into small experimental isolators and administrated orally with approximately 10^9^ CFUs (in 200 μL). Control animals remained in the breeding isolator during this period. After 11 days, all animals were exported into a laminar flow hood and sacrificed. Cecal content was collected for bile acid measurement and RNA extractions. RNA was extracted from 75-100 mg of cecal content obtained at the end of the experiment (11 days from the initiation of the colonisation experiment). 20 to 50 mg (dry weight) of cecal content were used for BA extraction. BA quantification and RT-qPCR assays were performed as described above.

### Statistical analysis and data visualisation

Graphpad Prism 9.2.0 (GraphPad) was used to generate the figures shown in this paper and perform pairwise comparisons, the gene expression data was analysed with a linear model (LM) in R language v4.1.2^62^ using RStudio^63^ a 2-way ANOVA or a Welch’s t-test. The statistical significance boundary was stablished at a *p*-value < 0.05.**Conflicts of interest**

### The authors declare no conflicts of interest

## Data Availability Statement

The data used for this manuscript are publicly available in the following link: https://doi.org/10.5281/zenodo.6034320

