## Supplementary Information for "Strain-dependent induction of primary bile acid 7-dehydroxylation by cholic acid"

*Bacterial growth in presence of bile acids un- and co-induced with $^{13}$C-CA.*

Growth was measured with $OD_{600}$ immediately before sample collection. All strains reached their maximum growth within 24 to 32 hours after the initial inoculation with an exponential phase culture after growth for approximately 12 hours (Supplementary Figure 10). The higher concentration of CDCA had an impact on bacterial growth although results were not as clear as previously reported[19]. Co-induction with $^{13}$C-CA did not have an impact on bacterial growth, as the co-induced groups generally followed the same growth curve as the CA group (without co-induction) (Supplementary Figure 10, D and F). The higher count of timepoints in the *C. scindens* ATCC 35704 was caused by a longer lag phase than expected which forced to follow closer the growth of this bacterium in order to capture the optimal RNA sample collection point (mid-log to late-log phase).

*In vitro transformation of βMCA and impact of $^{13}$C-CA co-induction*

As with αMCA, novel unknown compounds were detected in the presence of $^{13}$C-CA, albeit at lower concentrations and only in the highly active *C. scindens* VPI 12708. None of the strains tested had any detectable activity from βMCA when uninduced. *C. scindens* ATCC 35704 had minute amounts of an unidentified oxo form labelled X-oxoβMCA with a maximum of 3.38 µM at 32 hours (Supplementary Figure 9A) (hypothetically, 3-oxo-6β,7β-Dihydroxy-5β-cholan-24-oic acid). Co-inducing *C. scindens* VPI 12708 yielded a higher diversity of secondary BAs, although concentrations remained low. Small concentrations of 6-oxoMDCA were detected at the 32- and 48-hour time points, with its maximum at 0.42 µM after 32 hours (Supplementary Figure 9B). X- and Y- oxoβMCA forms were detected at maximum concentrations of 3.55 µM (32 hours) and 0.33 µM (48 hours), respectively (hypothetically, 3-oxo-6β,7β-Dihydroxy-5β-cholan-24-oic acid or 6-oxo-3α,7β-Dihydroxy-5β-cholan-24-oic acid). Moreover, very low concentrations of another BA with the same mass as 6-oxoMDCA (one oxidation and one dehydroxylation) were also detected at a maximum of 0.71 µM after 48 hours) (Supplementary Figure 9B) (currently unknown). *E. muris* SJ24 did not present any activity in the co-induced experiment (Supplementary Figure 9C), replicating the results observed from the αMCA group.

**Supplementary Table 1. Experimental layout for *in vitro* 7-dehydroxylation assays. Uninduced and co-induced experiments were performed at different times for technical reasons. CA (100 µM) was included in the co-induced experiment as a comparison to the uninduced experiment. Growth conditions can be found in Materials and Methods. No BA control included an equivalent volume of solvent (ethanol). All three bacteria were tested using the same design. Each group had biological triplicates.**

| Uninduced experiment | | | | | |
|---|---|---|---|---|---|
| 100 µM CA | 200 µM CDCA | 100 µM UDCA | 100 µM αMCA | 100 µM βMCA | No BA control |
| $^{13}$C-CA co-induced experiment | | | | | |
| 100 µM CA | 200 µM CDCA + 100 µM $^{13}$C-CA | 100 µM UDCA + 100 µM $^{13}$C-CA | 100 µM αMCA + 100 µM $^{13}$C-CA | 100 µM βMCA + 100 µM $^{13}$C-CA | No BA control |

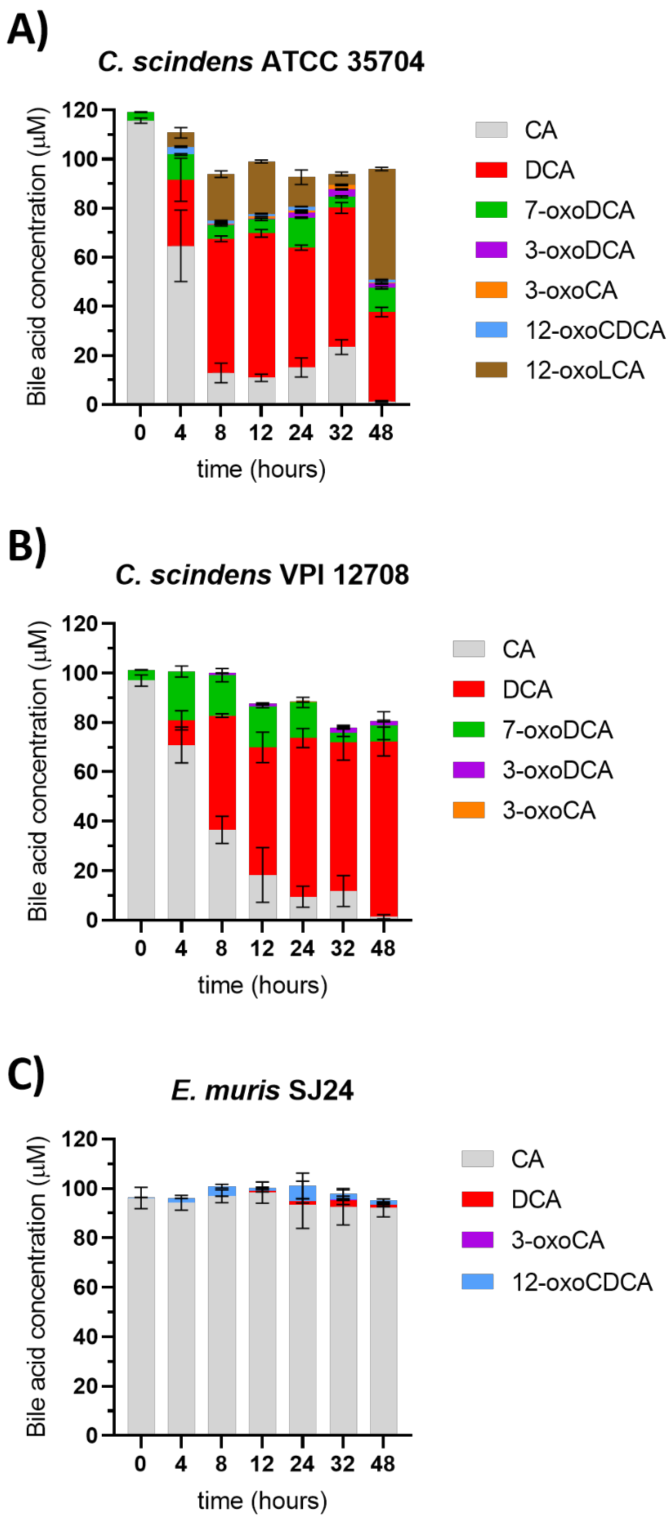


**Supplementary Figure 1.** ***In vitro*** **7-dehydroxylation of cholic acid by multiple strains. Bacteria were grown in BHIS-S in the presence of 100 µM of CA. This experiment is a repeat of that presented in Figure 2. However, it was performed at the same time as the experiment in which the three strains were co-induced with $^{13}$C-CA while the former was conducted at the same time as the experiment using uninduced strains (A)** ***Clostridium scindens*** **ATCC 35704, (B)** ***C. scindens*** **VPI 12708 and (C)** ***Extibacter muris*** **DSM 28561 (SJ24). Bile acids were measured post-extraction from the suspended biomass. Error bars represent the standard deviation of the mean of biological triplicates.**

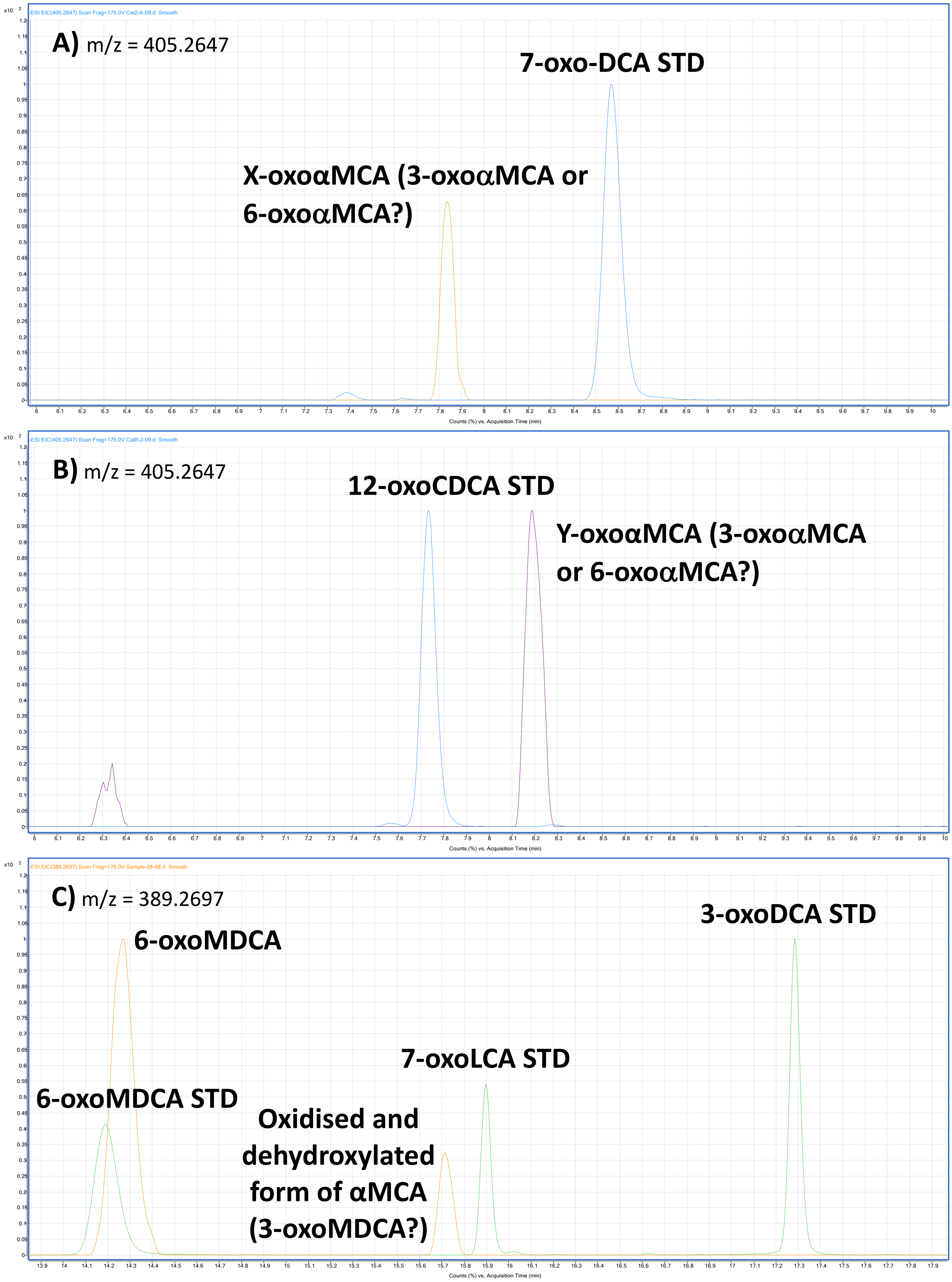
-ESI EIC(405.2647) Scan Frag=175.0V Cal2-A-09.d Smooth
A) m/z = 405.2647
7-oxo-DCA STD
X-oxoαMCA (3-oxoαMCA or 6-oxoαMCA?)
Counts (%) vs. Acquisition Time (min)
-ESI EIC(405.2647) Scan Frag=175.0V CalB-2-09.d Smooth
B) m/z = 405.2647
12-oxoCDCA STD
Y-oxoαMCA (3-oxoαMCA or 6-oxoαMCA?)
Counts (%) vs. Acquisition Time (min)
-ESI EIC(389.2697) Scan Frag=175.0V Sample-28-08.d Smooth
C) m/z = 389.2697
3-oxoDCA STD
6-oxoMDCA
7-oxoLCA STD
6-oxoMDCA STD
Oxidised and dehydroxylated form of αMCA (3-oxoMDCA?)
Counts (%) vs. Acquisition Time (min)

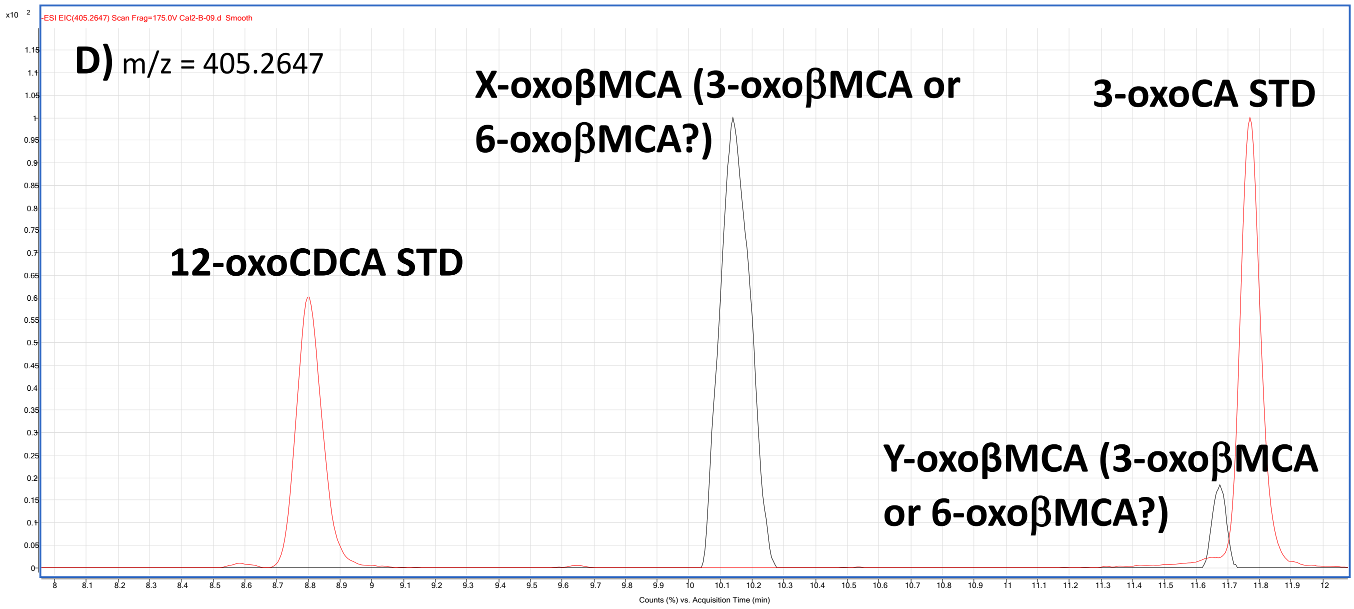
-ESI EIC(405.2647) Scan Frag=175.0V Cal2-B-09.d Smooth
D) m/z = 405.2647
X-oxoβMCA (3-oxoβMCA or 6-oxoβMCA?)
3-oxoCA STD
12-oxoCDCA STD
Y-oxoβMCA (3-oxoβMCA or 6-oxoβMCA?)
Counts (%) vs. Acquisition Time (min)

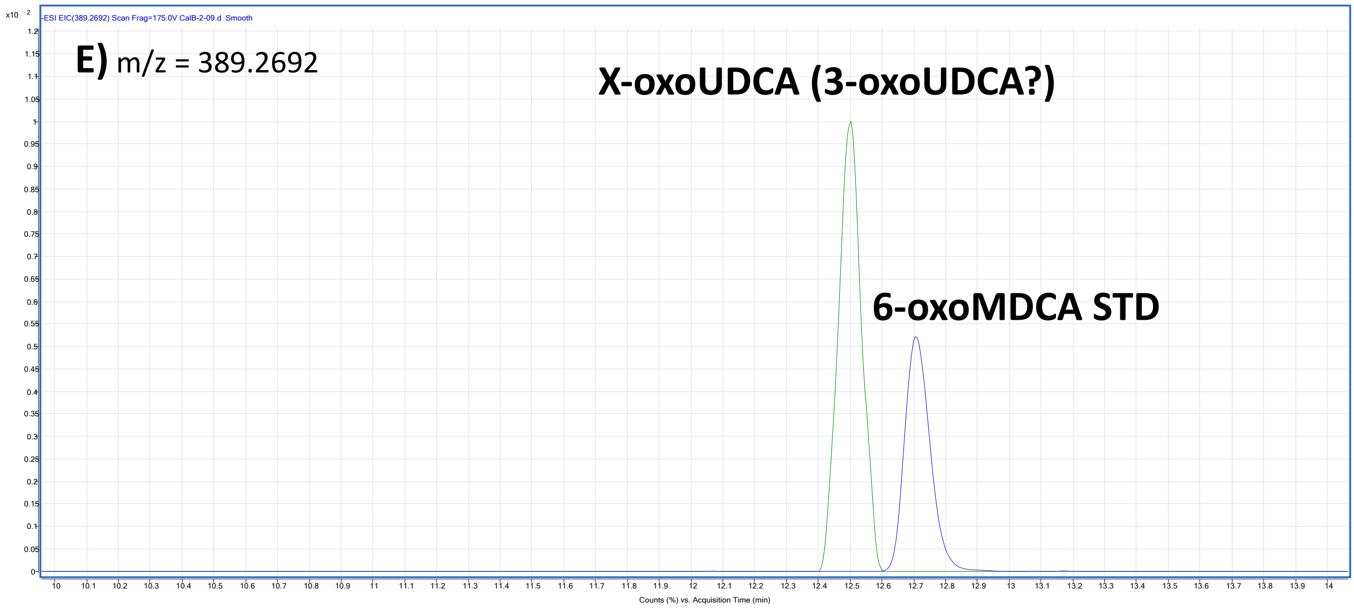
-ESI EIC(389.2692) Scan Frag=175.0V CalB-2-09.d Smooth
E) m/z = 389.2692
X-oxoUDCA (3-oxoUDCA?)
6-oxoMDCA STD
Counts (%) vs. Acquisition Time (min)

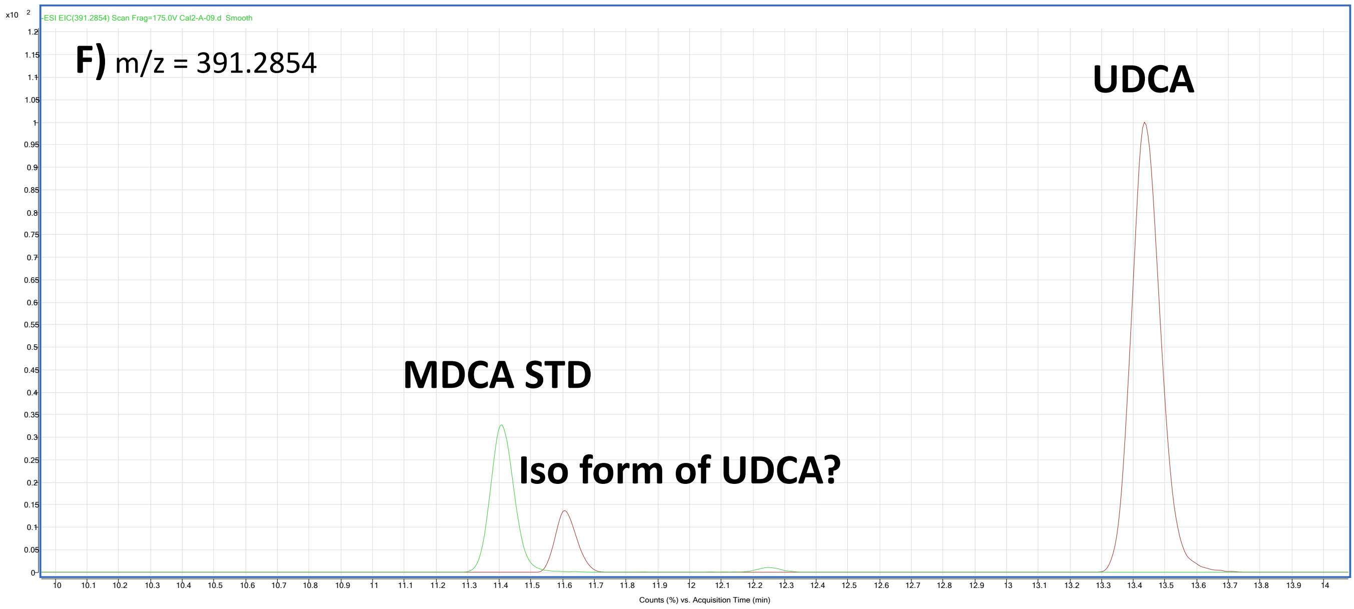


**Supplementary Figure 2. Extracted chromatograms for unknown bile acids identified in this study. Extracted chromatograms were chosen from a representative sample which contained one or more of the unknown BAs. Y axis represents signal intensity, X axis represents acquisition time in minutes (retention time). Chromatograms were set to "scale to largest in each chromatogram". Shortened names are written in bold on top or near the peak. STD indicates standard. Extracted mass for each chromatogram is indicated on the top left corner. (A) X-oxoαMCA suggested to be 3- or 6- oxo-αMCA: 3-oxo-6β,7α-Dihydroxy-5β-cholan-24-oic acid or 6-oxo-3α,7α-Dihydroxy-5β-cholan-24-oic acid respectively. (B) Y-oxoαMCA suggested to be 3- or 6- oxo-αMCA: 3-oxo-6β,7α-Dihydroxy-5β-cholan-24-oic acid or 6-oxo-3α,7α-Dihydroxy-5β-cholan-24-oic acid respectively. We can't assign a specific compound to either X-oxoαMCA or Y-oxoαMCA but can state that one is 3-oxo-αMCA and the other 6-oxo-αMCA. (C) The product of the oxidation and dehydroxylation of αMCA, suggested to be 3-oxoMDCA (3-oxo-6β-Hydroxy-5β-cholan-24-oic acid). (D) X-oxoβMCA and Y-oxoβMCA suggested to be 3- or 6- oxo-βMCA; 3-oxo-6β,7β-Dihydroxy-5β-cholan-24-oic acid or 6-oxo-3α,7β-Dihydroxy-5β-cholan-24-oic acid respectively. We can't assign a specific compound to either X-oxoβMCA or Y-oxoβMCA but can state that one is 3-oxo-βMCA and the other 6-oxo-βMCA. (E) X-oxoUDCA suggested to be 3-oxoUDCA (3-oxo-7β-Hydroxy-5β-cholan-24-oic acid). (F) Proposed iso form of UDCA (3β,7β-Dihydroxy-5β-cholan-24-oic acid).**

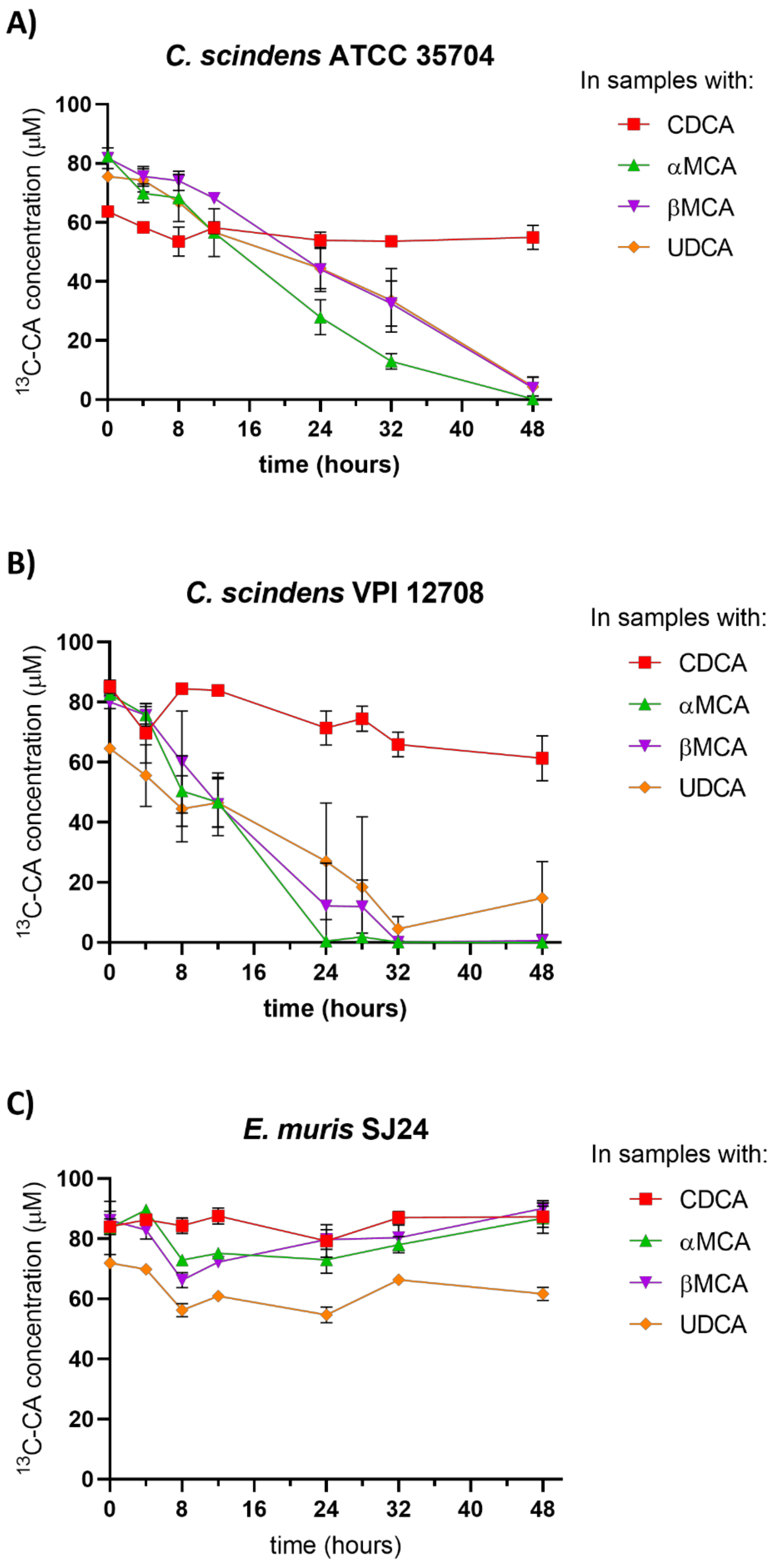


**Supplementary Figure 3. 7-dehydroxylation of $^{13}$C-CA in the presence of the various primary bile acids tested (CDCA, αMCA, βMCA, UDCA). (A) *Clostridium scindens* ATCC 35704, (B) *C. scindens* VPI 12708 and (C) *Extibacter muris* DSM 28561 (SJ24). 100 µM of $^{13}$C-CA was used to induce 7-dehydroxylation of the other primary bile acids. Error bars represent the standard deviation of the mean from triplicates.**

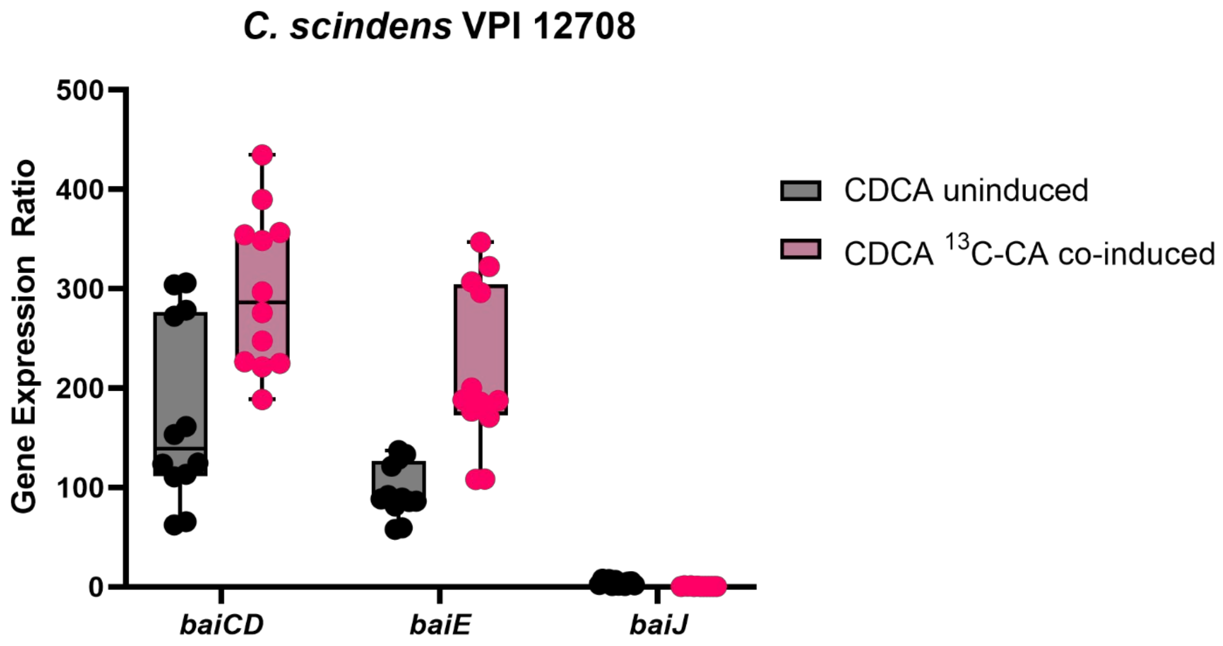


**Supplementary Figure 4.** ***C. scindens*** **VPI 12708** ***baiCD*****,** ***baiE*** **and** ***baiJ*** **gene expression ratio in CDCA with and without** $^{13}$**C-CA co-induction. This figure is a detail of the same values shown in Figure 6B. Coloured dots represent individual replicates. Boxplot shown with minimum and maximum bars. Differences between uninduced and co-induced conditions were significant (**) with a** ***p*****-value < 0.01 when using a paired Wilcoxon test but not when doing a more complete linear model analysis as performed in Figure 6.**

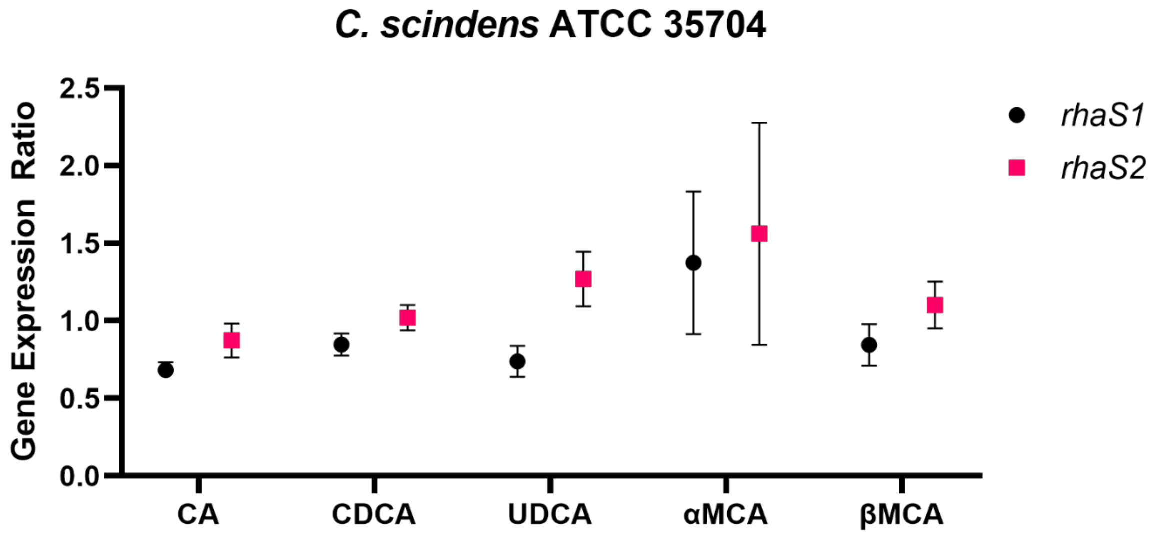


**Supplementary Figure 5.** ***rhaS*** **gene expression in the presence of various primary BAs in** ***Clostridium scindens*** **ATCC 35704. gene expression of** ***rhaS1 and rhaS2*****. The** ***rhaS1*** **gene is also known as** ***barA*****. At least three reference genes were used and a control group with an equivalent volume of solvent (ethanol) to that used in the groups with bile acids. CA, UDCA, αMCA, βMCA were used at 100 µM, CDCA was used at 200 µM. Dots represent the average and error bars represent the standard deviation from 12 replicates.**

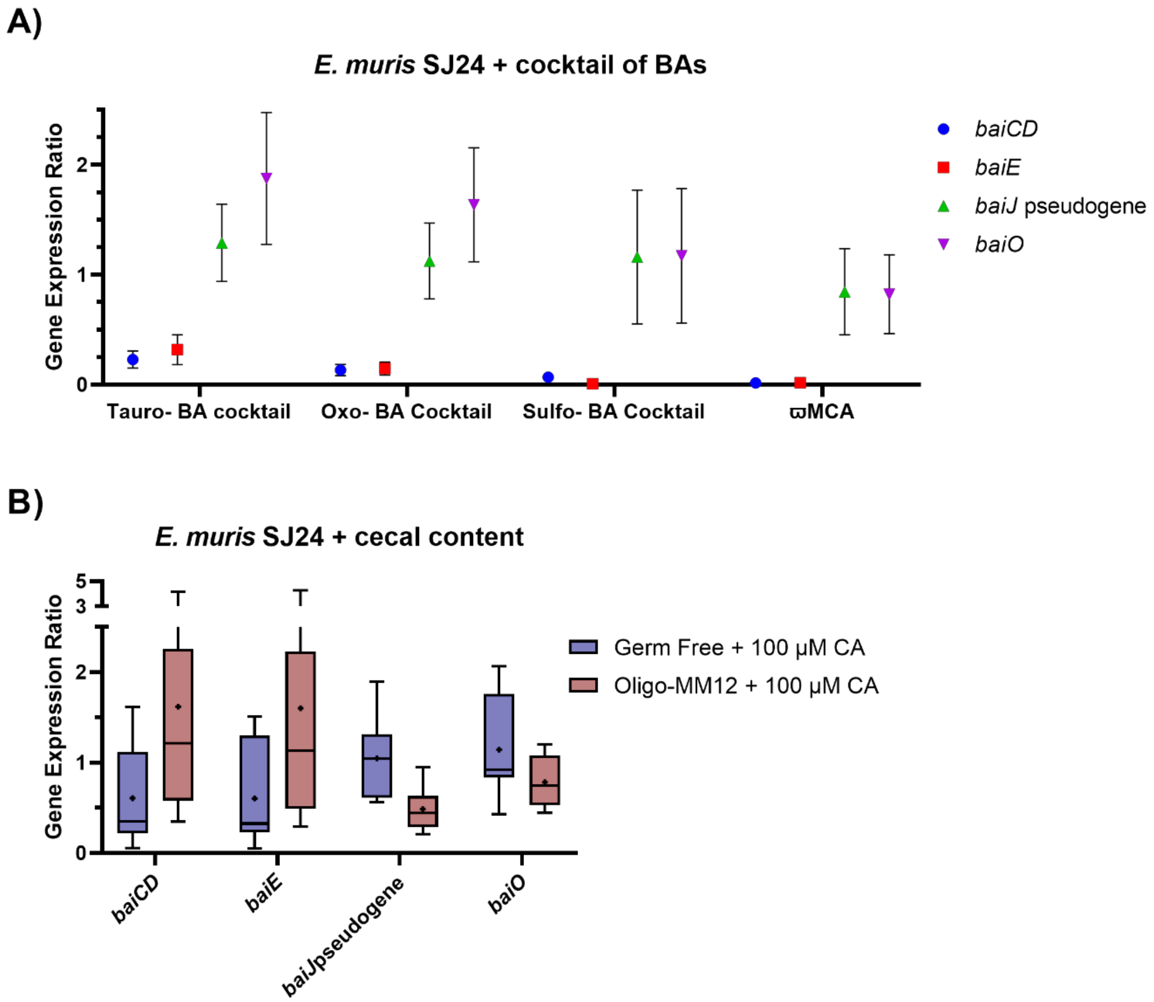


**Supplementary Figure 6.** ***E. muris*** **DSM 28561 (SJ24)** ***baiCD*****,** ***baiE*****, pseudogene** ***baiJ*** **and** ***baiO*** **gene expression. (A) In presence of various BA cocktails and (B) in presence of 25 mg of cecal content from germ-free mice or from oligo-MM12 mice (cecal content mixed with 5% glycerol). For each experiment, the expression (normalised to at least three reference genes) is relative to a 100 µM CA condition that was included as a control. The composition of the BA cocktails is detailed in the materials and methods section of the paper. (A) Coloured dots represent the average and error bars represent the standard deviation of 12 replicates. (B) Box and whiskers highlight five-number summary for each group (maximum value, 75th percentile, median, 25th percentile and minimum value), the dot within each box shows the position of the mean. A two-way ANOVA did not find any significance in the differences between treatment groups for either assay.**

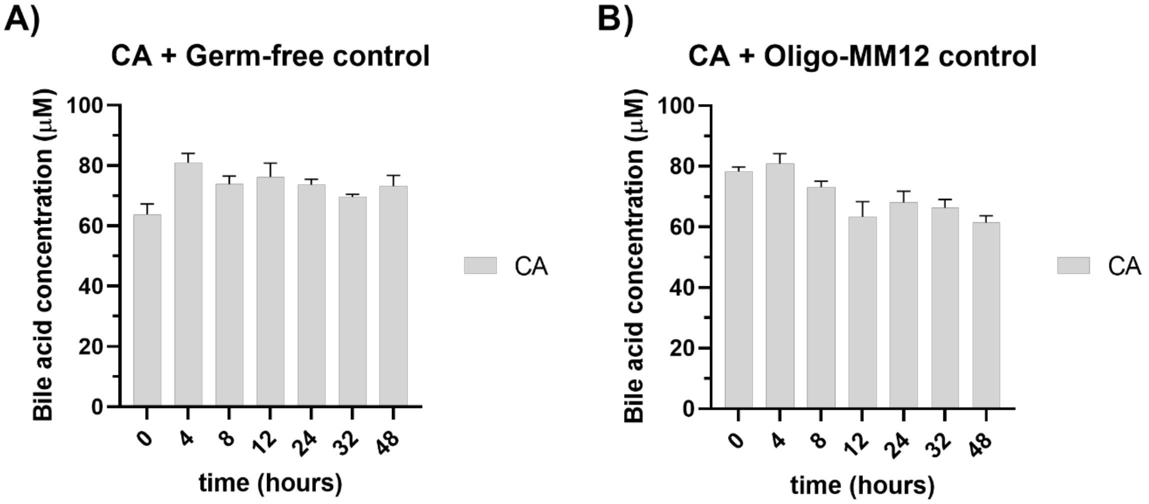


**Supplementary Figure 7. Control groups of cecal content with 100 µM of CA added. Control groups from Figure 8. 25 mg of (A) cecal content from germ-free mice and (B) cecal content from Oligo-MM12 colonized mice. The mass imbalance between the added CA (100 µM) and the measured (aprox. 80 µM) can be attributed to the presence of CoA- forms that cannot currently be quantified due to lack of pure standards. Error bars represent the standard deviation of biological triplicates.**

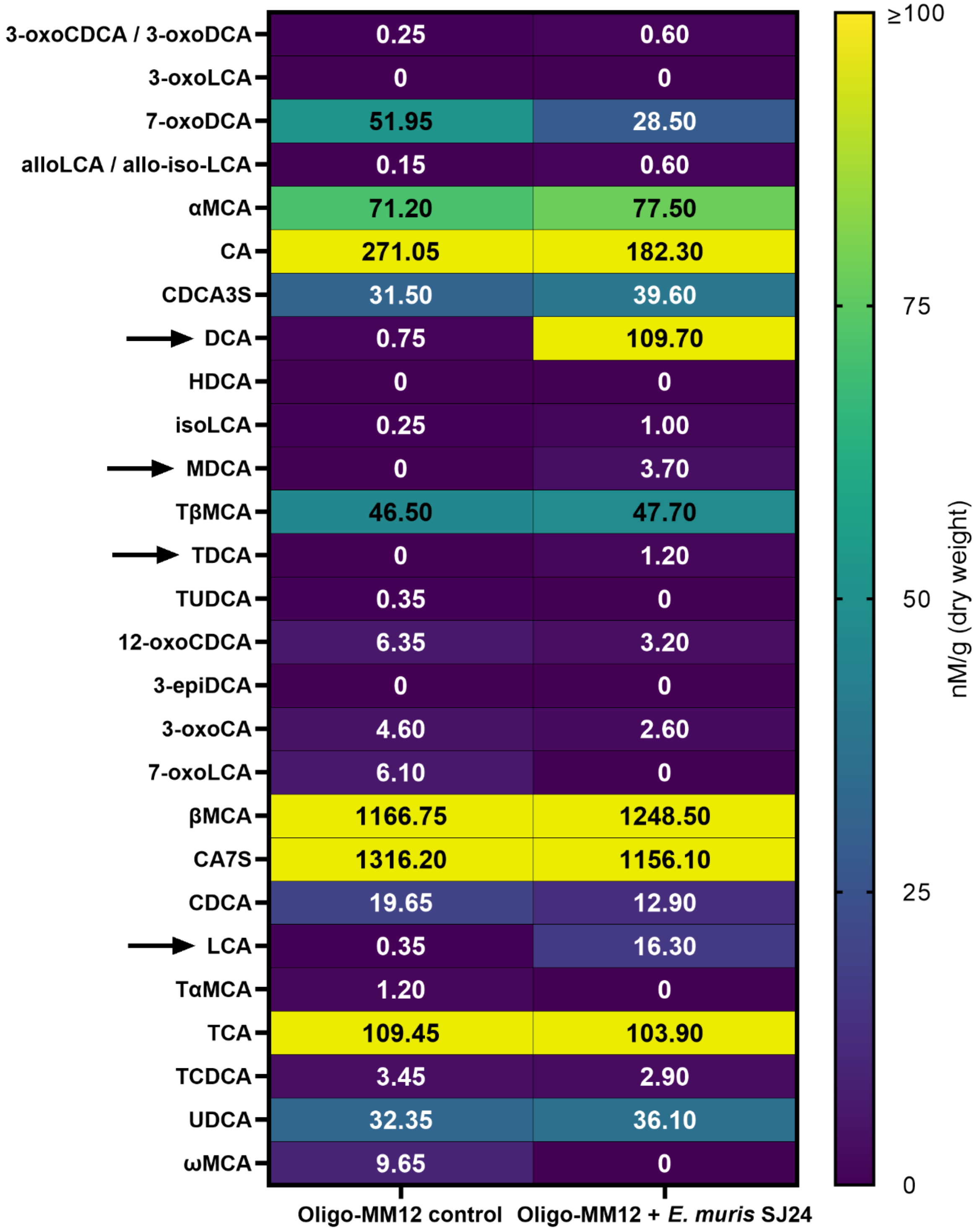


**Supplementary Figure 8. BA pool of Oligo-MM12 mice with and without *E. muris* SJ24. BAs were extracted from cecal content frozen immediately after sacrifice and until extraction and quantification. Results are presented as nM per gram of dry weight. Values within the cells represent the median value of 4 (Oligo-MM12 control) or 5 (Oligo-MM12 + *E. muris* SJ24) biological replicates. Arrows are used to highlight the secondary 7-DH-ed BAs DCA, TDCA, LCA and MDCA, exclusively present in the Oligo-MM12 + *E. muris* SJ24 group.**

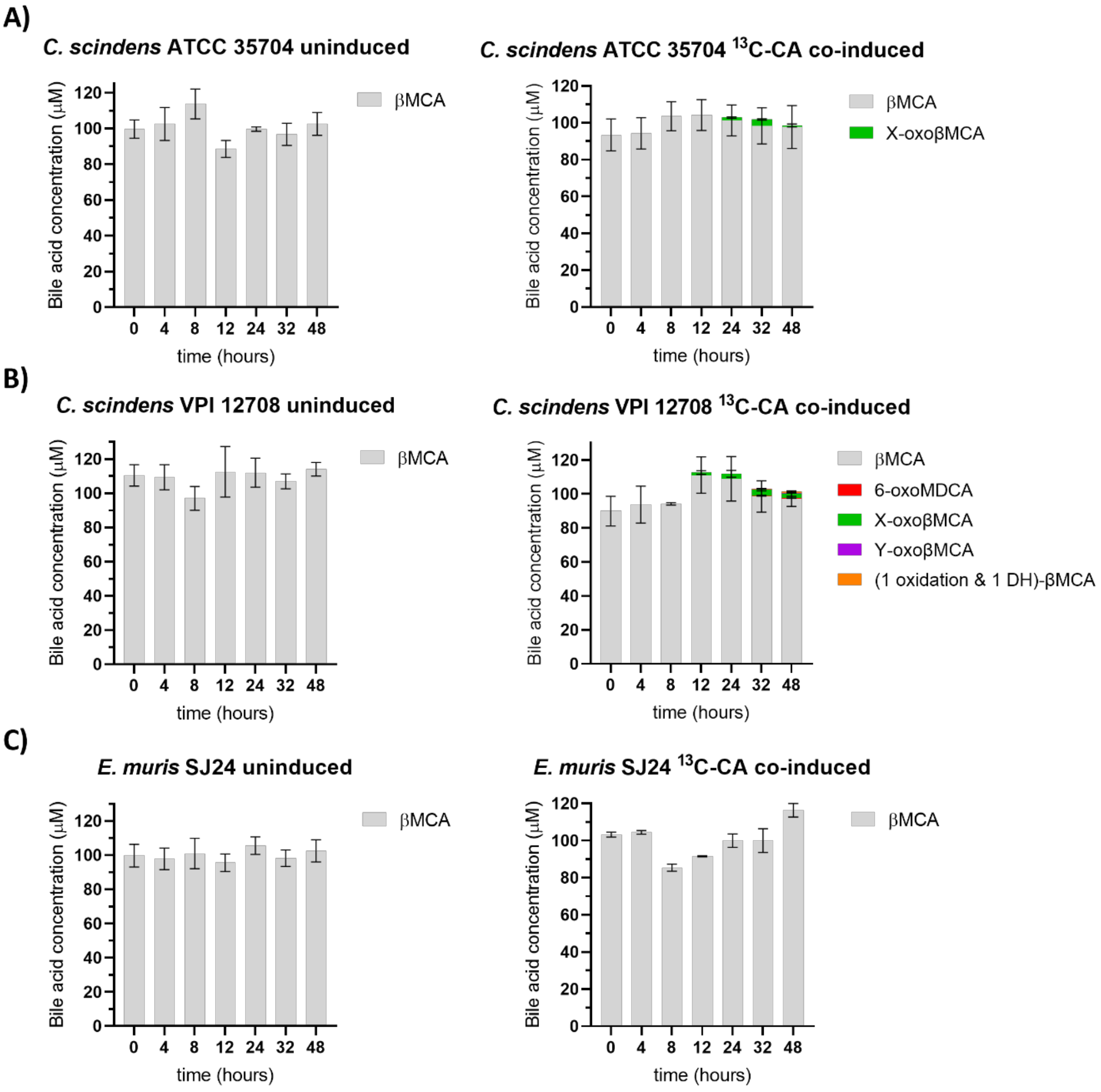


**Supplementary Figure 9.** ***In vitro*** **7-dehydroxylation of βMCA. The transformation of 100 µM of βMCA into secondary bile acids was tested with and without co-induction with 100 µM of $^{13}$C-CA. (A)** ***Clostridium scindens*** **ATCC 35704, (B)** ***C. scindens*** **VPI 12708 and (C)** ***Extibacter muris*** **DSM 28561 (SJ24) were grown anaerobically in BHIS-S. Bile acids were extracted from suspended biomass. Several compounds were detected that could not be identified due to missing standards but their oxidative state can be estimated based on their ionised mass. X- or Y- oxoβMCA had the same mass as other bile acids with one ketone group and two hydroxyl groups. The other unidentified compound had the same mass as secondary bile acids that have been dehydroxylated (-1 -OH), have one ketone group and one hydroxyl group. The retention times for these compounds was unique and therefore could not be identified further. Concentration values of the unknown BAs could only be estimated for the same reason. Error bars represent the standard deviation of the mean of biological triplicates.**

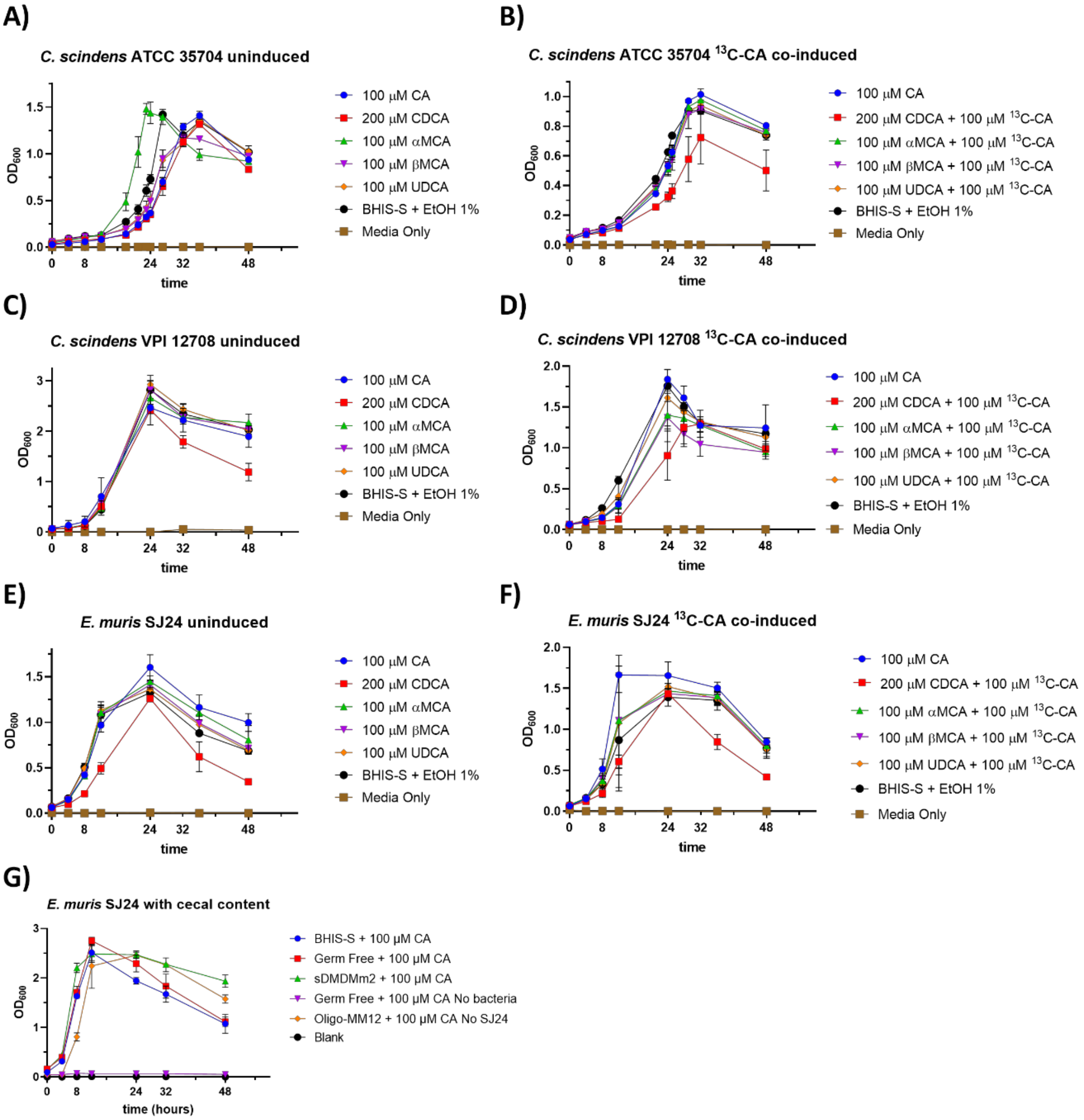


**Supplementary Figure 10. Growth curves of 7-dehydroxylating bacteria strains in various primary bile acids. Bacteria were grown anaerobically in BHIS-S medium with CA, CDCA, αMCA, βMCA, or UDCA (A-F) or with 25 mg of cecal content (G) and a control with only the ethanol solvent (BHIS-S + EtOH 1%). The co-induced groups (B, D and F) received an additional 100 µM $^{13}$C-CA. A sterile control was included (Media Only). (A-B) *Clostridium scindens* ATCC 35704, (C-D) *C. scindens* VPI 12708, (E-G) *Extibacter muris* DSM 28561 (SJ24). Error bars represent the standard deviation calculated from biological triplicates.**

**Supplementary Table 2. Primer table. Primers labelled "Scindens" were used for both *Clostridium scindens* ATCC 35704 and VPI 12708 strain. All primers were tested for efficiency and redesigned if necessary.**

| Primer name | Primer sequence 5' – 3' |
|---|---|
| Scindens_recA.Fw | GGAGAACGCCAAGTCATATCTG |
| Scindens_recA.Rv | TCTTCCGACTTCTCTTCTCCTG |
| Scindens_rho.Fw | AAGATATGACCCCGATCTTCCC |

| | |
|---|---|
| Scindens_rho.Rv | GGATACGATCATTCCCCTCTGT |
| Scindens_rpsJ.Fw | GTAGTGACAATCTTAAGGGCGG |
| Scindens_rpsJ.Rv | CAGCCGGCATCTCTAATCTAGA |
| Scindens_adk.Fw | GCCGCAAAGTATGGTATTCCTC |
| Scindens_adk.Rv | GTACCAGAAGTCCCTGATCCAT |
| Scindens_16S.Fw | TATTAGGAGGAACACCAGTGGC |
| Scindens_16S.Rv | CTAGTAGTCATCGTTTACGGCG |
| Scindens_gyrB.Fw | CCGGATGACACTATTTTCGAGG |
| Scindens_gyrB.Rv | CTCCGGTCTTTCATCCCTAAGT |
| Scindens_rpoB.Fw | TCCACAGGACCATATTCTCTGG |
| Scindens_rpoB.Rv | GACTTCACAGTCAGGATCTCCT |
| Scindens35704_baiCD.Fw | CTTAAGAACCGTATCGTCCTGC |
| Scindens35704_baiCD.Rv | CCGGACATAAGGCTACACATTC |
| Scindens35704_baiE.Fw | GATCTTTACGGACGGCAAGTAC |
| Scindens35704_baiE.Rv | GCCTGTTTCAAGGATGTACCAC |
| Scindens35704_baiJ.Fw | GCAGAGAGAGCAGGGAATCTAT |
| Scindens35704_baiJ.Rv | CCATCTTGATCAGCCATTCCAG |
| Scindens12708_baiCD.Fw | GAATACCGAAGTGACTGCAGAG |
| Scindens12708_baiCD.Rv | CATAGGTAGAGCCTACTGCGAT |
| Scindens12708_baiE.Fw | AAAGACGTAGGAATCAATGGCG |
| Scindens12708_baiE.Rv | GGATCTTTGGATCACGCATGAA |
| Scindens12708_baiJ.Fw | CCAGTATCATGTATCCGATGCG |
| Scindens12708_baiJ.Fw | CTGCTTGAACCCATACTGTACG |
| MurisSJ24_L3p.Fw | AAGATGCCTGGACATATGGGAA |
| MurisSJ24_L3p.Rv | CTAAAGATTTCTTCGGTCCCGG |
| MurisSJ24_16S.Fw | GGGGATAACAGCTAGAAATGGC |
| MurisSJ24_16.Rv | TACCGCACCAACTACCTAATCA |
| MurisSJ24_recA.Fw | GGGAGAAGGTAGGGATCATGTT |
| MurisSJ24_recA.Rv | ACGGATAGACGCGTAGAACTTA |
| MurisSJ24_rpoB.Fw | AGGTATGGAAGAAAAAGCTGCC |

| | |
|---|---|
| MurisSJ24_rpoB.Rv | CTCATTGGAAGTAGAGCGTTCC |
| MurisSJ24_gyrB.Fw | CGGTTGAAAAATCAGTCATGGC |
| MurisSJ24_gyrB.Rv | TTTTTGGGTCTTTGTCGGAACA |
| MurisSJ24_baiCD.Fw | TGTGACTCCGGAACTGATAGAG |
| MurisSJ24_baiCD.Rv | CCTTTGAGCACTTCATACTGGG |
| MurisSJ24_baiE.Fw | ATCGTGACCTCTTACTCCGATG |
| MurisSJ24_baiE.Rv | GCGGTAGTATCGCTGTCTATTG |
| MurisSJ24_pseudogenebaiJ.Fw | GGCAGAAGAAACAAGGCATCTT |
| MurisSJ24_pseudogenebaiJ.Rv | CGTCTTTGCACGTATAGAAGCT |
| MurisSJ24_baiO.Fw | ATTCTATCGAGACGATGGGAGG |
| MurisSJ24_baiO.Rv | CTCCTGTCTGCGTATCTTCAAC |

**Supplementary Table 3. List of bile acids quantified with the LC-MS. (*) Also known as 6-oxolithocholic acid (6-oxoLCA)**

| Bile acid common name | Abbreviations | Synonym |
|---|---|---|
| 12-oxolithocholic acid | 12-oxoLCA | (3α,5β)-3-Hydroxy-12-oxocholan-24-oic acid |
| 3-oxochenodeoxycholic acid | 3-oxoCDCA | 5β-Cholanic acid-12α-ol-3-one |
| 3-oxolithocholic acid | 3-oxoLCA | 3-Oxo-5β-cholan-24-oic acid |
| 6-oxo-allolithocholic acid | 6-oxo-alloLCA | 5α-Cholanic acid-3α-ol-6-one |
| 7-oxodeoxycholic acid | 7-oxoDCA | 7-Ketodeoxycholic acid |
| Allolithocholic acid | alloLCA | (5α)-3β-Hydroxy-cholan-24-oic acid |
| α-muricholic acid | αMCA | 5β-Cholanic acid-3α,6β,7α-triol |
| Cholic acid | CA | 3α,7α,12α-Trihydroxy-5β-cholanic acid |
| Chenodeoxycholic acid-3-sulfate | CDCA3S | (3α,5β,7α)-7-Hydroxy-3-(sulphooxy)-cholan-24-oic Acid |
| Deoxycholic acid | DCA | 3α,12α-Dihydroxy-5β-cholanic acid sodium salt |
| Hyocholic acid | HCA | 3α,6α,7α-Trihydroxy-5β-cholanic acid |
| Hyodeoxycholic acid | HDCA | 3α,6α-Dihydroxy-5β-cholan-24-oic acid |
| Isolithocholic acid | isoLCA | (3β,5β)-3-Hydroxy-cholan-24-oic acid |
| Murideoxycholic acid | MDCA | 5β-Cholanic acid-3α, 6β-Diol |
| Tauro-β-muricholic acid | TβMCA | 5β-Cholanic acid-3α, 6β, 7β, -triol N-(2-sulphoethyl)-amide |
| Taurocholic acid-3-sulfate | TCA3S | Taurocholic acid 3-sulfate |

| | | |
|---|---|---|
| Taurodeoxycholic acid | TDCA | Taurodeoxycholate |
| Taurolithocholic acid-3-sulfate | TLCA3S | Taurolithocholic acid 3-sulfate |
| Tauroursodeoxycholic acid | TUDCA | 3α,7β-Dihydroxy-5β-cholan-24-oic acid N-(2-sulphoethyl)-amide |
| Tauro-ω-muricholic acid | TωMCA | 5β-Cholanic acid-3α,6α,7β,-triol N-(2-sulphoethyl)-amide |
| 12-oxoChenodeoxycholic acid | 12-oxoCDCA | 3α,7α-Dihydroxy-12-oxo-5β-cholanic acid |
| 24-$^{13}$C-cholic acid | 24-$^{13}$C-CA | Cholic-24-$^{13}$C acid |
| 3-epi-deoxycholic acid | 3-epiDCA | 5β-Cholanic acid-3β,12α-diol |
| 3-oxocholic acid | 3-oxoCA | 3-Oxo-7α,12α-hydroxy-5β-cholanoic acid |
| 3-oxodeoxycholic acid | 3-oxoDCA | 5β-Cholanic acid-12α-ol-3-one |
| 6-oxomurideoxycholic acid* | 6-oxoMDCA | 5-β-Cholanic acid-3α-ol-6-one |
| 7,12-dioxolithocholic acid | 7,12-dioxoLCA | 5β-Cholanic acid-3α-ol-7, 12-dione |
| 7-oxolithocholic acid | 7-oxoLCA | 3α-Hydroxy-7-oxo-5β-cholanic acid |
| β-muricholic acid | βMCA | 5β-Cholanic acid-3α,6β,7β-triol |
| Cholic acid-7-sulfate | CA7S | (3α,5β,7α,12α)-3,12-Dihydroxy-7-(sulfooxy)-cholan-24-oic acid |
| Chenodeoxycholic acid | CDCA | 5β-Cholanic acid-3α,7α-diol |
| Isodeoxycholic acid | isoDCA | 5β-Cholanic acid-7α,12α-diol |
| Lithocholic acid | LCA | 3α-Hydroxy-5β-cholan-24-oic acid |
| Tauro-α-muricholic acid | TαMCA | 5β-Cholanic acid-3α, 6β, 7α, -triol N-(2-sulphoethyl)-amide |
| Taurocholic acid | TCA | Taurocholate |
| Taurochenodeoxycholic acid | TCDCA | Taurochenodeoxycholate |
| Taurohyocholic acid | THCA | 5β-Cholanic acid-3α, 6α, 7α-triol N-(2-sulphoethyl)-amide |
| Taurohyodeoxycholic acid | THDCA | Taurohyodeoxycholate |
| Taurolithocholic acid | TLCA | Taurolithocholic acid |
| Tauromurideoxycholic acid | TMDCA | Tauromurideoxycholic acid |
| Ursodeoxycholic acid | UDCA | 5β-Cholan-24-oic acid-3α,7β-diol |
| ω-muricholic acid | ωMCA | 5β-Cholanic acid-3α, 6α, 7β-triol |
